# Engineering tumor-homing bacteria as membrane-anchored immune checkpoint-blockade interfaces

**DOI:** 10.64898/2026.06.10.731477

**Authors:** Zongqi Wang, Chengyuan Fang, Xingwu Zhou, Shaobo Yang, Yatrik M. Shah, Rizwan Romee, James J. Moon, Jiahe Li

## Abstract

Bacterial cancer therapies can exploit tumor tropism to localize immunomodulatory payloads, but most approaches rely on secretion or lysis-dependent release of soluble biologics that may diffuse beyond the tumor niche. Here, we engineer non-pathogenic, tumor-homing *Escherichia coli* strains as membrane-anchored immunotherapeutic interfaces, with individual strains displaying immune checkpoint-blocking nanobodies targeting CTLA-4 or PD-L1, either alone or in combination with a separate strain displaying murine decoy-resistant IL-18 (mDR18). By screening multiple bacterial outer-membrane proteins as scaffolds, we identified scaffold-dependent differences in display levels and target engagement, and YiaT as a functional platform for checkpoint nanobody presentation. Delivery of YiaT-displayed nanobodies in combination with OmpA-displayed mDR18 suppressed tumor growth in syngeneic mouse colon cancer and melanoma models in either local or systemic delivery. Upon systemic administration, the combined bacterial therapy preferentially accumulated in tumors, outperformed benchmark checkpoint antibody regimens combining CTLA-4 and PD-L1 under the tested conditions, and promoted tumor rejection and rechallenge resistance, without inducing broad systemic cytokine release. Further immune profiling showed that the combined treatment was associated with increased CD8⁺ and effector-memory T-cell responses in tumors and spleens. This work establishes bacterial surface display as a modular strategy for localized cancer immunotherapy.

## Introduction

Standard cancer treatment options such as chemotherapy and radiotherapy often show limited efficacy in solid tumors, particularly in the setting of poor vascularization, hypoxia, and immune suppression ^1^. Microbial-based cancer therapy represents a promising strategy to overcome the limitations of conventional cancer treatment while synergizing with existing immunotherapy ^2–6^. This is because certain bacterial strains preferentially accumulate in hypoxic and necrotic regions of solid tumors, which are difficult to access with conventional biologics ^6^. They can either directly activate the immune system to attack tumor cells or can be engineered to deliver a myriad of therapeutic payloads ^2,5,7^. Notably, several genera, including *Escherichia*, *Salmonella*, *Listeria*, *Clostridium,* and *Bifidobacterium*, have been explored as tumor-targeting therapeutic platforms ^8–10^, and some bacterial products have advanced into clinical testing ^7,8,11–16^. As self-renewing living therapeutics, microbial platforms may also offer practical advantages for scalable and potentially lower-cost cancer treatment.

Previous microbial cancer therapies have primarily relied on secretion or programmed lysis to release soluble immunotherapeutic payloads, including nanobody-based checkpoint blockers, after tumor colonization ^10,17–23^. Although these approaches have established the feasibility of bacterial delivery of immune-modulatory biologics, soluble payloads may exhibit poor pharmacokinetics, potentially limiting local retention and increasing systemic exposure ^24^. Consistent with this concern, broader local immunotherapy studies have shown that anchoring potent cytokines to tumor matrices, such as genetic fusion of immune cytokines with collagen-binding peptides or proteins, can improve local retention, safety, and efficacy ^24–26^. However, matrix-anchoring strategies are generally optimized for local administration and may not fully address the challenge of systemically delivering immune agonists to inaccessible tumor sites.

Toward this goal, our group recently established a bacterial surface-display platform in which non-pathogenic, tumor-homing *Escherichia coli* are armored with membrane-bound immune cytokines. In this study, surface-displayed murine decoy-resistant IL-18 (DR18) enhanced antitumor immunity, synergized with immune checkpoint blockade, and potentiated T cell- and NK cell-mediated tumor control ^27^. These findings are consistent with broader efforts in engineering adoptively transferred T cells or NK cells, in which membrane-bound cytokines can retain efficacy while reducing systemic exposure ^28,29^. Thus, previous work by others and our own effort support the idea that anchoring potent immune agonists to cell surfaces can concentrate their activities at the therapeutic site while reducing the toxicity associated with systemic cytokine exposure. In the present work, we reasoned that bacterial surface display could extend this anchoring principle to immune checkpoint blockade by converting tumor-homing bacteria from passive payload carriers into membrane-anchored checkpoint-blockade interfaces. In this setting, immune checkpoint blockers are presented directly on the bacterial outer membrane, potentially increasing local avidity, restricting activity to sites of bacterial accumulation, and enabling modular combination with additional surface-displayed immune agonists.

We chose to focus on PD-L1 as an especially attractive target for bacteria-localized immune checkpoint blockade because its immunosuppressive function can arise from both tumor and host compartments ^30^. Prior work comparing syngeneic mouse tumor models demonstrated that PD-L1 on MC38 colorectal tumor cells is sufficient to suppress antitumor immunity and directly inhibit CD8⁺ T-cell cytotoxicity, whereas PD-L1 on non-tumor host cells plays a major role in limiting antitumor immunity in B16F10 melanoma ^30^. Thus, targeting PD-L1 locally within tumors may help intercept inhibitory PD-1/PD-L1 interactions across malignant and tumor-associated host-cell compartments, providing a rationale for evaluating surface-displayed PD-L1 blockade in both colorectal cancer and melanoma models. In parallel, CTLA-4 blockade provides a complementary immune checkpoint target by modulating T-cell priming and costimulatory signaling ^10,31^, supporting combined local targeting of CTLA-4 and PD-L1 pathways.

Therefore, we engineer non-pathogenic, tumor-homing *E. coli* strains that individually display immune checkpoint-blocking nanobodies targeting PD-L1 or CTLA-4, or combine these single-payload strains with a separate strain that displays mDR18 ^27,32^. Through systematically characterizing outer-membrane proteins that support the functional presentation of immune checkpoint nanobodies, we demonstrate scaffold-dependent antitumor efficacy after intratumoral delivery. Using the optimized constructs, we further show that systemic administration of the combined bacterial therapy suppresses tumor growth, promotes durable immune memory, and avoids broad systemic induction of inflammatory cytokines. These results establish bacterial surface display as a versatile strategy for enabling localized, membrane-anchored cancer immunotherapies that integrate immune checkpoint blockade and cytokine stimulation at tumor sites.

## Results

### Outer-membrane protein scaffold engineering identifies YiaT as an optimal interface for functional immune checkpoint nanobody display

We began by screening bacterial outer-membrane protein (OMP) scaffolds for their ability to display checkpoint nanobodies while preserving target engagement. We selected the non-pathogenic facultative anaerobe *E. coli* K-12 as the bacterial chassis because of its reported tumor tropism, ease of genetic manipulation, and pan-sensitivity to commonly used antibiotics ^33^. We performed a head-to-head comparison of five commonly used bacterial OMP constructs: outer membrane protein A (OmpA), putative outer membrane protein YiaT truncated at 181^st^ amino acid (YiaT181), N-terminal EaeA protein (Neae), autotransporter EhaA C-terminal fragment (C-EhaA), and adhesin involved in diffuse adherence (AIDA) (**Figure 1a and Extended Data Fig. 1**). In these constructs, αCTLA-4 (PDB: 5E03) or αPD-L1 (PDB: 5DXW) nanobodies ^10,34^ were fused to the C terminus of OmpA ^35^, YiaT181 ^36^, and Neae ^37^, or to the N terminus of the autotransporter scaffolds C-EhaA ^38^ and AIDA ^39^, according to the expected extracellular domain of each OMP. The expression of each surface display construct was detected using an anti-nanobody antibody for flow cytometry. Among the scaffolds tested, Neae resulted in the highest overall surface-display signal for both nanobodies, followed by YiaT181, whereas C-EhaA and AIDA showed comparatively lower display levels (**Figure 1b**). Because C-EhaA and AIDA require N-terminal nanobody fusion, whereas OmpA, YiaT181, and Neae use C-terminal fusion, the reduced display from C-EhaA and AIDA may reflect orientation-dependent effects on folding, translocation, or surface accessibility.

**Figure 1.**
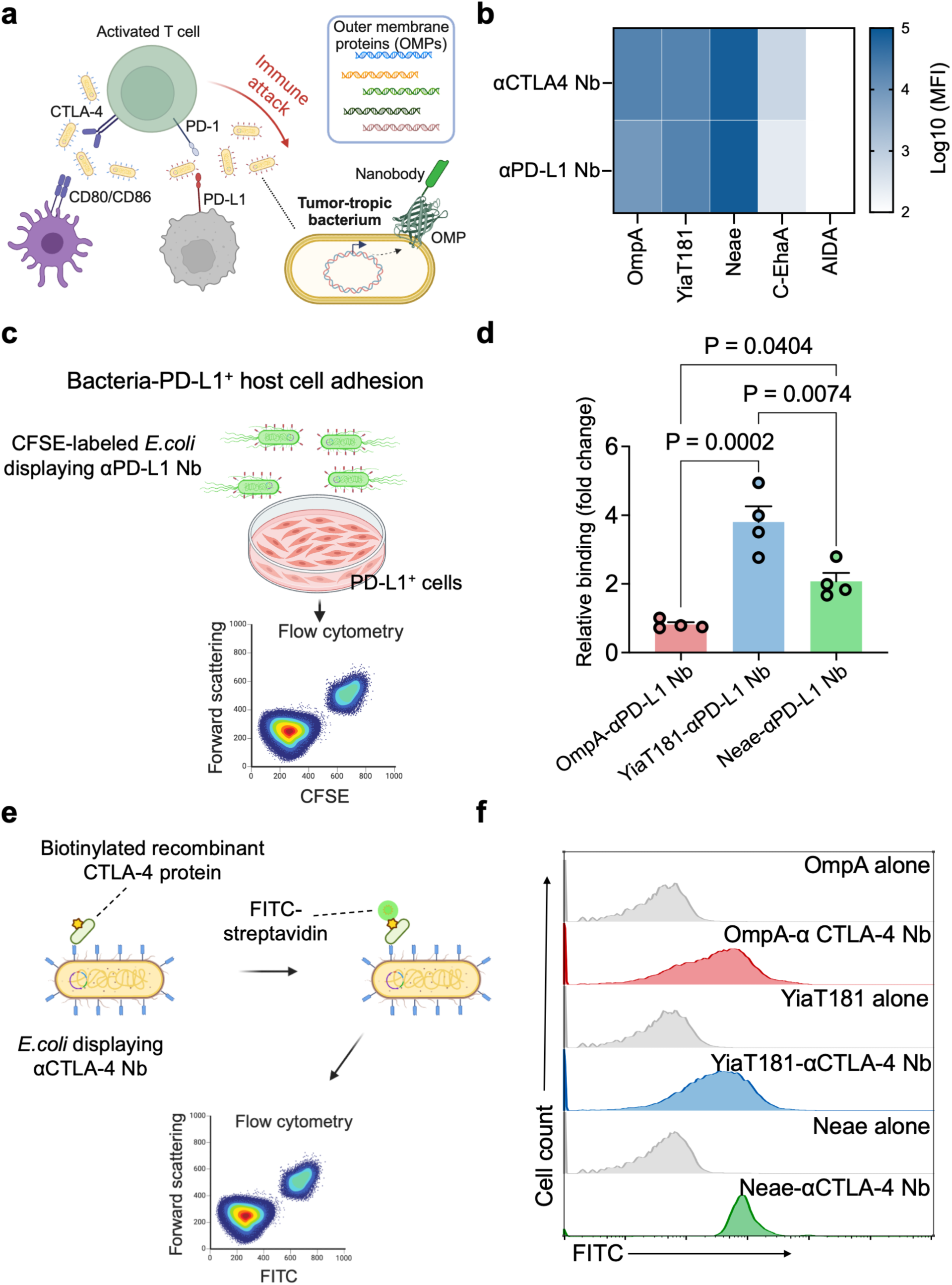
Engineering and characterization of bacterial surface display platforms for immune checkpoint nanobody presentation. **a,** Schematic illustration of engineered *E. coli* displaying αCTLA-4 or αPD-L1 nanobody (Nb) to modulate inhibitory immune checkpoint interactions. **b,** Heatmap showing surface display levels of αCTLA-4 and αPD-L1 nanobodies using different outer membrane display scaffolds. The C terminus of outer membrane protein A (OmpA), putative outer membrane protein truncated at 181^st^ amino acid (YiaT181), or N-terminal EaeA protein (Neae) was fused to the nanobody of interest, while the N terminus of autotransporter EhaA C-terminal fragment (C-EhaA) or adhesin involved in diffuse adherence (AIDA) was fused to the nanobody. The display levels were detected with an anti-nanobody antibody, measured by flow cytometry, and reported as log10-transformed mean fluorescence intensity (MFI). Data are representative of two independent biological replicates. **c,** Schematic of the bacteria-host cell binding assay. Engineered *E. coli* displaying αPD-L1 nanobodies were fluorescently labeled with CFSE and incubated with mouse DC 2.4 dendritic cells at a multiplicity of infection (MOI) of 100, and analyzed by flow cytometry. **d,** Quantification of bacterial binding to DC 2.4 cells for the indicated αPD-L1 nanobody display scaffolds. Relative binding was determined by flow cytometry and normalized to the corresponding scaffold-only control strain (OmpA-αPD-L1 to OmpA, YiaT181-αPD-L1 to YiaT181, and Neae-αPD-L1 to Neae). Each dot represents an individual sample (n = 4 technical repeats from one of the two independent biological replicates); bars indicate mean ± s.e.m. Statistical significance was determined by one-way ANOVA followed by Tukey’s multiple-comparisons test. Exact P values are indicated in the figure. Data are representative of two independent biological experiments. **e,** Schematic of the recombinant CTLA-4 binding assay. Engineered *E. coli* displaying αCTLA-4 nanobodies were incubated with biotinylated recombinant CTLA-4 protein, followed by FITC-conjugated streptavidin staining and flow cytometric analysis. **f,** Representative flow cytometry histograms showing FITC fluorescence following incubation of bacteria displaying αCTLA-4 nanobodies with biotinylated recombinant CTLA-4 protein and FITC-conjugated streptavidin. Corresponding scaffold-only controls are shown in gray. Data are representative of two independent biological replicates.

We next asked whether the displayed nanobodies retained functional target-binding activity. For PD-L1 targeting, engineered *E. coli* was first stained with carboxyfluorescein succinimidyl ester (CFSE) and then incubated with mouse dendritic cells constitutively expressing PD-L1 ^40^. αPD-L1-dependent bacterial adhesion to dendritic cells was measured by flow cytometry, while *E. coli* engineered to display empty scaffold proteins served as negative controls (**Figure 1c**). YiaT181-mediated display induced the strongest binding to DC 2.4 cells, whereas Neae-supported display showed detectable but lower binding activity (**Figure 1d**). In contrast, OmpA-displayed αPD-L1 showed limited binding compared with the OmpA-only control (**Extended Data Fig. 2a**) . These results indicate that high surface-display level alone is not sufficient to predict functional target engagement. Similar results were obtained when B16F10 melanoma cells were used as another PD-L1-positive cell line (**Extended Data Fig. 2b**).

To evaluate CTLA-4 targeting, we incubated engineered bacteria displaying αCTLA-4 nanobodies with biotinylated recombinant mouse CTLA-4 protein, then stained with FITC-conjugated streptavidin and analyzed by flow cytometry (**Figure 1e**). In contrast to the more OMP-dependent pattern for αPD-L1 binding, OmpA-, YiaT181-, and Neae-mediated αCTLA-4 display showed comparable CTLA-4 binding relative to their scaffold-only controls, indicating that each OMP scaffold supported functional CTLA-4 binding (**Figure 1f**). These findings show that the choice of outer-membrane scaffold influences both nanobody display and target engagement, with the optimal scaffold varying across nanobody-target pairs.

### Membrane-anchored immune checkpoint nanobody display confers scaffold-dependent antitumor activity after intratumoral delivery

In addition to the *in vitro* OMP screening, we further determined whether these surface-displayed immune checkpoint nanobodies could translate into antitumor activity *in vivo*. Prior work showed that the MC38 model is sensitive to immune checkpoint blockade ^27,41^. We therefore used MC38 as an initial testbed to compare surface-display scaffolds in a local delivery setting, where differences in scaffold-dependent nanobody function could be directly evaluated (**Figure 2a**). C57BL/6J mice bearing subcutaneous MC38 tumors (∼100 mm^3^) received intratumoral doses of engineered bacterial strains separately displaying individual αCTLA-4 or αPD-L1 nanobodies (Nbs) on distinct outer-membrane scaffolds. Hereafter, “Nbs” refers to a mixture of two single-payload strains separately displaying αCTLA-4 or αPD-L1. We prioritized combined CTLA-4 and PD-L1 blockade over testing each checkpoint blocker because our goal was to compare scaffold-dependent display performance and maximize the likelihood of detecting scaffold-dependent *in vivo* activity in this initial proof-of-concept study. Consistent with the weak αPD-L1 binding observed for the OmpA-based construct *in vitro*, OmpA-Nbs did not show a clear therapeutic benefit compared with the OmpA scaffold-only control on day 18 (**Figure 2b**). By contrast, YiaT181-Nbs induced the strongest tumor growth suppression compared with the corresponding YiaT181-only control, whereas Neae-Nbs showed a more modest antitumor effect relative to Neae alone (**Figure 2b**). Longitudinal tumor monitoring further supported the superior *in vivo* performance of the YiaT181-based construct, which delayed tumor progression and extended survival relative to the other scaffold configurations in the MC38 model (**Figure 2c, d**). In summary, these data indicate that the OMP scaffold choice influences not only nanobody display and target binding *in vitro*, but also therapeutic efficacy *in vivo*. Therefore, we prioritized YiaT181-Nbs and Neae-Nbs for subsequent *in vivo* combination studies and did not further test OmpA-Nbs.

**Figure 2.**
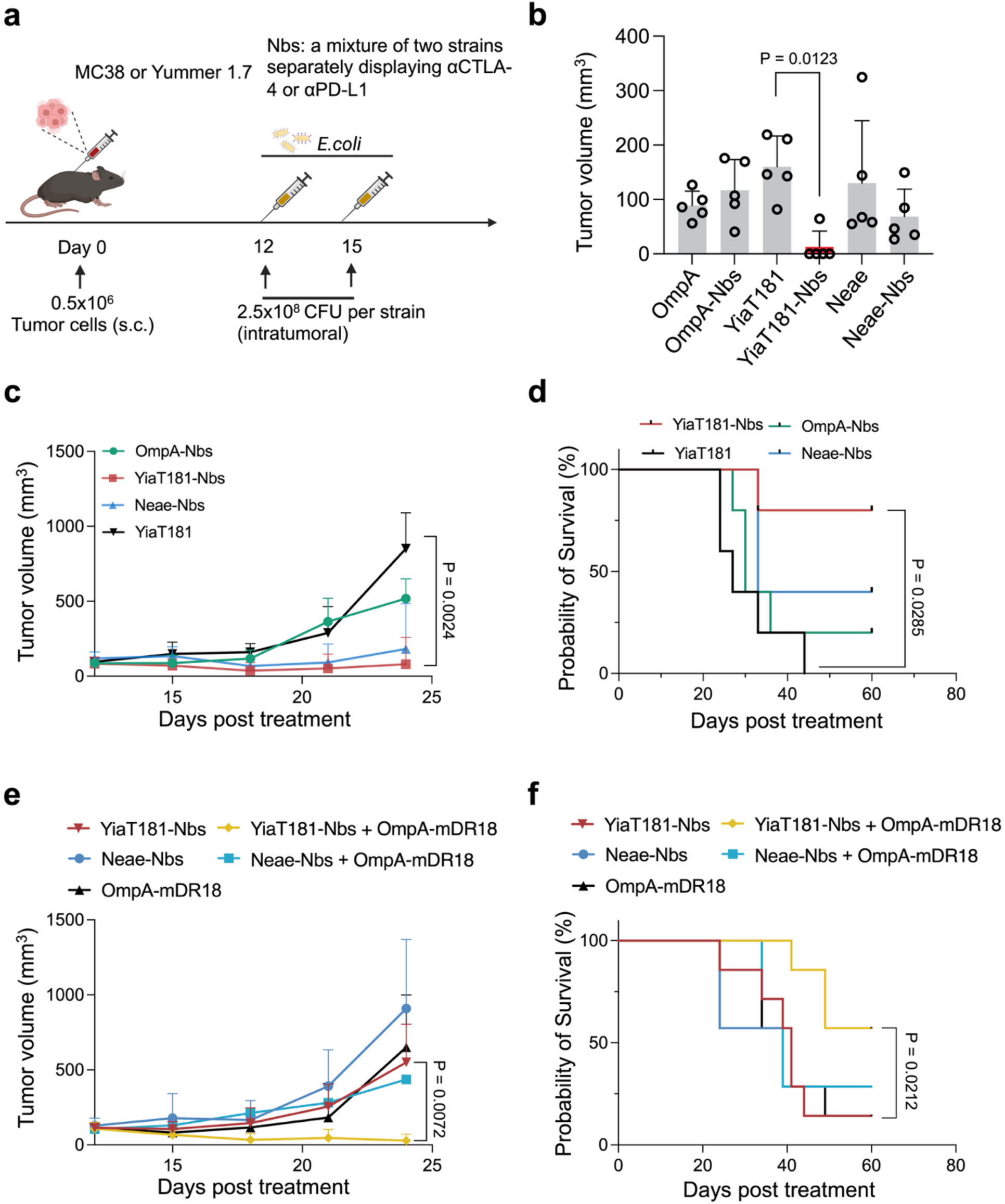
Evaluation of bacterial surface display scaffolds and intratumoral therapeutic efficacy of checkpoint nanobody-displaying bacteria. **a,** Schematic illustration of the intratumoral treatment regimen. MC38 or Yummer1.7 tumor-bearing mice were established by subcutaneous inoculation of tumor cells on day 0 and treated intratumorally with engineered *E. coli* at the indicated time points. Throughout this figure, “Nbs” refers to a mixture of two single-payload strains separately displaying αCTLA-4 or αPD-L1. Per-strain bacterial doses were matched across treatment groups, with total CFU adjusted according to the number of co-administered strains as described in Methods. **b,** Tumor volumes measured on day 18 in MC38 tumor-bearing mice following treatment with bacteria expressing checkpoint nanobodies using different outer membrane display scaffolds. Each dot represents an individual mouse (n = 5 mice per group); bars indicate mean ± s.e.m. Statistical analysis was performed using one-way ANOVA followed by Tukey’s multiple-comparisons test. **c,** Tumor growth curves of MC38 tumor-bearing mice treated with OmpA-Nbs, YiaT181-Nbs, Neae-Nbs, or the corresponding scaffold-only control (YiaT181) (n = 5 mice per group). Data are presented as mean ± s.e.m. Statistical analysis was performed using two-way ANOVA followed by Tukey’s multiple-comparisons test. **d,** Kaplan-Meier survival curves corresponding to the treatment groups shown in **c (**n = 5 mice per group). Statistical significance was determined using the log-rank (Mantel-Cox) test. **e,** Tumor growth curves of Yummer1.7 tumor-bearing mice treated with checkpoint nanobody-displaying bacteria, OmpA-mDR18, or the indicated combination therapies (n = 7 mice per group). Data are presented as mean ± s.e.m. Statistical analysis was performed using two-way ANOVA followed by Tukey’s multiple-comparisons test. **f,** Kaplan-Meier survival curves corresponding to the treatment groups shown in **e (**n = 7 mice per group). Statistical significance was determined using the log-rank (Mantel-Cox) test.

Having identified YiaT181 as the strongest immune checkpoint nanobody-display scaffold in MC38, we next asked whether this module could be combined with a complementary membrane-anchored cytokine signal to enhance antitumor immunity. We evaluated immune checkpoint nanobody display, alone or in combination with OmpA-mDR18, in the Yummer1.7 melanoma model. Yummer1.7 is a syngeneic melanoma model carrying *Braf*^V600E^, *Pten*^−/−^, and *Cdkn2a*^−/−^ driver alterations, with relevance to human melanoma ^42^. Because this model is responsive to immune checkpoint therapy and supports evaluation of antitumor immune responses, it provided a suitable setting to test whether localized cytokine stimulation could enhance bacteria-based immune checkpoint nanobody therapy. Additionally, our previous work identified OmpA as the lead OMP scaffold for mDR18 surface display and antitumor activity, compared with YiaT and Neae ^27^. Following intratumoral administration of engineered *E. coli* in mice bearing established Yummer1.7 tumors, combining OmpA-mDR18 with YiaT181-Nbs resulted in the strongest suppression of tumor growth and improved survival, whereas the Neae-Nbs combination showed a less pronounced benefit (**Figure 2e, f**). In contrast, OmpA-mDR18 alone or checkpoint nanobody display alone was less effective. In summary, the intratumoral efficacy studies in two different syngeneic tumor models identified YiaT181 as the preferred scaffold for checkpoint nanobody display. They further supported combining YiaT181-displayed checkpoint nanobodies with OmpA-displayed mDR18 for subsequent systemic delivery studies.

### Systemic co-administration of bacteria separately displaying immune checkpoint nanobodies or mDR18 induces tumor control and durable immune memory

Having identified YiaT181 as the preferred scaffold for immune checkpoint nanobody display in combination with cytokine delivery, we next evaluated whether this optimized construct could mediate antitumor activity following systemic delivery. Yummer1.7-bearing mice received intravenous administration of engineered *E. coli* strains individually displaying YiaT181-anchored αCTLA-4 or αPD-L1 nanobodies, an OmpA-displayed mDR18 strain, or the combined bacterial treatment comprising these single-payload strains (**Figure 3a**). As benchmark controls, we included commercially available αCTLA-4 plus αPD-L1 blocking antibodies, which have been validated for *in vivo* antitumor efficacy, either alone or in combination with OmpA-mDR18 bacteria ^10,30,31^.

**Figure 3.**
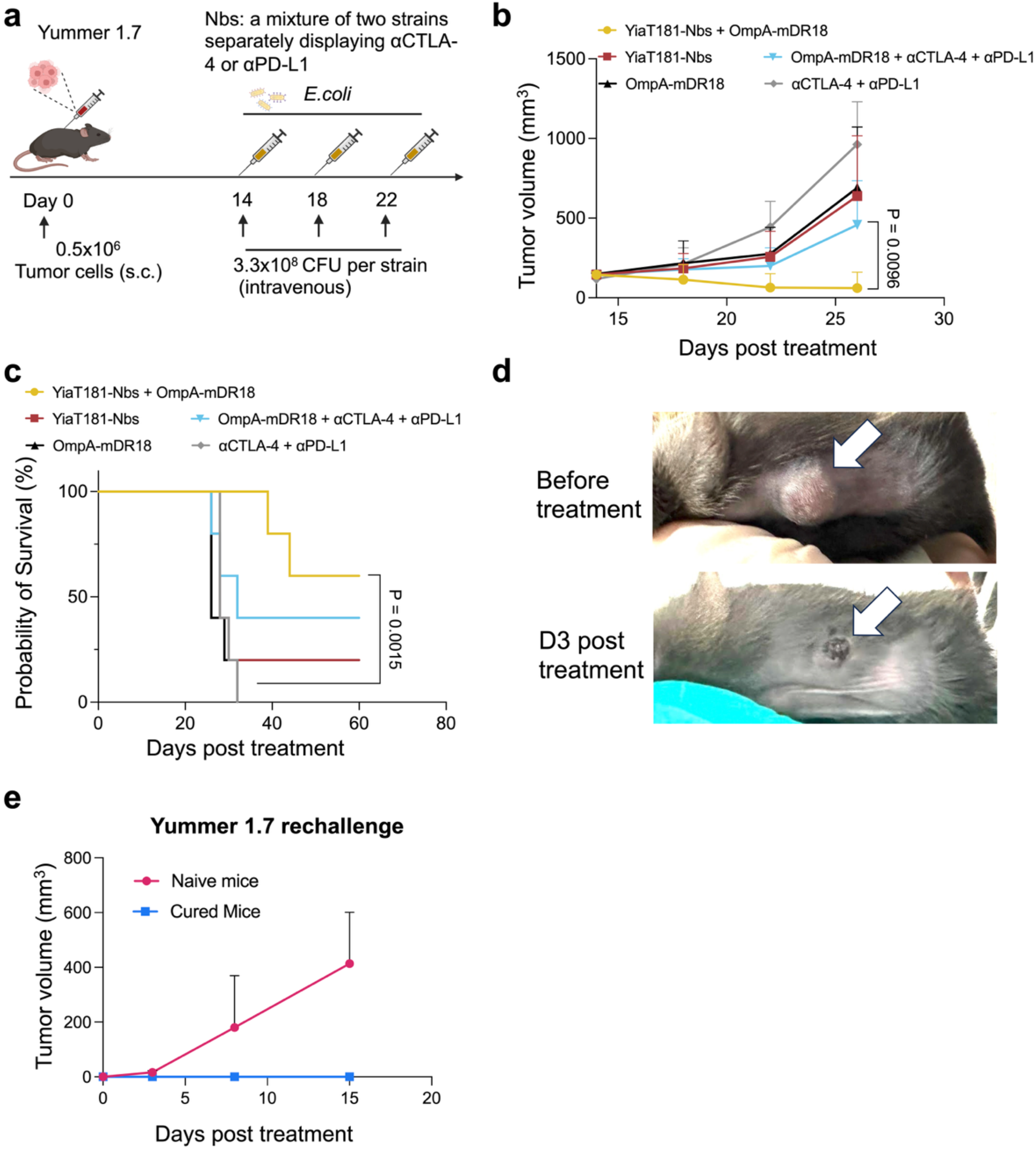
Systemic co-administration of engineered bacteria separately displaying checkpoint nanobodies or mDR18 mediates antitumor activity and immunological memory. **a,** Schematic illustration of the treatment regimen in the Yummer1.7 melanoma model. Mice were subcutaneously inoculated with Yummer1.7 tumor cells on day 0 and subsequently treated intravenously with engineered *E. coli* at the indicated time points. Throughout this figure, “Nbs” refers to a mixture of two single-payload strains separately displaying αCTLA-4 or αPD-L1. Bacterial treatment groups received matched per-strain doses, with total CFU adjusted according to the number of co-administered strains as described in Methods. Antibody-treated groups received 200 μg anti-CTLA-4 (clone 9D9) and 100 μg anti-PD-L1 (clone 10F.9G2) per mouse. **b,** Tumor growth curves of Yummer1.7 tumor-bearing mice treated with YiaT181-Nbs, OmpA-mDR18, YiaT181-Nbs plus OmpA-mDR18, OmpA-mDR18 combined with commercial checkpoint blockade antibodies, or commercial checkpoint blockade antibodies alone (n = 5 mice per group). Data are presented as mean ± s.e.m. Statistical analysis was performed using two-way ANOVA followed by Tukey’s multiple-comparisons test. The combination of commercial αCTLA-4 (clone 9D9) + αPD-L1 (clone 10F.9G2) blocking antibodies serves as a benchmark. **c,** Kaplan-Meier survival analysis of mice receiving the treatments shown in **b** (n = 5 mice per group). Statistical significance was determined using the log-rank (Mantel-Cox) test. **d**, Representative images of Yummer1.7 tumors before treatment and 3 days after treatment with co-administered YiaT181-Nbs and OmpA-mDR18 bacteria. **e**, Tumor rechallenge study evaluating the development of immunological memory. Mice that completely rejected primary tumors following treatment were rechallenged with Yummer1.7 cells and compared with sex- and age-matched naïve mice (n = 3 mice per group). Tumor growth was monitored after rechallenging.

Among the groups tested, co-administration of YiaT181-Nbs with OmpA-mDR18 induced the strongest suppression of tumor growth and the most favorable survival outcome (**Figure 3b, c**). In comparison, systemic treatment with commercial αCTLA-4 and αPD-L1 blocking antibodies, alone or combined with OmpA-mDR18 bacteria, was less effective under these experimental conditions. These results indicate that intravenous administration preserved the activity observed after intratumoral delivery, supporting the feasibility of systemic bacterial delivery for localized tumor immunotherapy. Furthermore, combining membrane-anchored checkpoint blockade with localized cytokine stimulation enhances therapeutic efficacy. To determine whether this treatment could induce durable antitumor immunity, we rechallenged mice that had completely rejected Yummer1.7 tumors after bacterial therapy. Notably, cured mice resisted tumor re-engraftment, whereas age- and sex-matched naïve controls developed progressive tumors (**Figure 3d, e**). These findings suggest that treatment with engineered bacteria not only mediates initial tumor clearance but also elicits protective immunological memory.

We next asked whether this therapeutic strategy could extend to a more aggressive B16F10 melanoma model. In this model, checkpoint blockade alone is typically less effective ^43,44^, and prior studies have implicated non-tumor host-cell PD-L1 as an important suppressor of antitumor immunity in B16F10 ^30^. This allowed us to test whether systemically delivered bacteria could present αPD-L1 nanobodies within a tumor microenvironment in which PD-L1-mediated suppression is not restricted to malignant cells. Intravenous delivery of a combination of engineered *E. coli* strains separately displaying YiaT181-anchored αCTLA-4, YiaT181-anchored αPD-L1, or OmpA-mDR18 produced the greatest therapeutic benefit, with the strongest reduction in tumor growth relative to bacteria displaying either component alone or commercial checkpoint blockade (**Extended Data Fig. 3**). In summary, these findings demonstrate that systemically delivered engineered *E. coli* can localize to tumors and that the combination of YiaT181-displayed checkpoint nanobodies with OmpA-displayed mDR18 provides an effective membrane-anchored immunotherapy approach across distinct syngeneic tumor settings.

### Tumor-enriched bacterial localization is associated with limited systemic inflammatory toxicity after intravenous delivery

Because systemic administration of live bacteria and membrane-bound immunostimulatory payloads raises important biodistribution and safety considerations, we next evaluated where engineered *E. coli* accumulated after intravenous delivery. This analysis was particularly important because the therapeutic concept depends on preferential bacterial enrichment in tumors, where checkpoint nanobodies and mDR18 are intended to act locally, rather than in healthy organs, where off-tumor immune activation could increase toxicity. We focused on tumors, spleens, livers, and kidneys because these tissues are associated with the major biological and safety questions for systemically delivered bacterial immunotherapy. Tumors represent the intended therapeutic site, where bacterial enrichment would support localized presentation of checkpoint nanobodies and mDR18. The spleen and liver are major reticuloendothelial organs involved in systemic immune surveillance and bacterial clearance, and therefore provide sensitive readouts for off-tumor bacterial accumulation and inflammatory risk. The kidney was included to assess potential bacterial dissemination to a highly perfused organ relevant to systemic toxicity and tissue clearance.

Three days after intravenous administration, engineered *E. coli* showed strong tumor-selective accumulation, with approximately 1,000- to 10,000-fold higher bacterial burden in tumors than in liver, spleen, or kidney tissues (**Figure 4a**). This preferential tumor enrichment is consistent with the known ability of facultative anaerobic bacteria to colonize hypoxic and necrotic regions within solid tumors, and supports the concept that bacterial surface display can localize immunostimulatory biologics to the tumor microenvironment while limiting exposure in major healthy organs. We next asked whether this tumor-biased localization translated into a safe systemic profile. Mice receiving intravenous bacterial therapy showed no obvious adverse effects, including no overt weight loss (**Figure 4b**) or hunched posture (data not shown). To evaluate systemic inflammatory responses, we analyzed plasma or tumor lysate samples collected 7 days after bacterial administration using a 32-plex Luminex panel. Compared with PBS-treated controls, mice receiving engineered bacterial therapies did not show elevation of cytokines and chemokines commonly associated with cytokine-release syndrome, such as IL-6, IL-1β, TNFα, IFNγ, GM-CSF, MCP-1/CCL2, IP-10/CXCL10, and G-CSF (**Figure 4c, Extended Data Figures 5** and **6**). Therefore, the bacterial biodistribution and cytokine profiling support the potential safety of using bacterial surface display to concentrate checkpoint blockade and cytokine within tumors.

**Figure 4.**
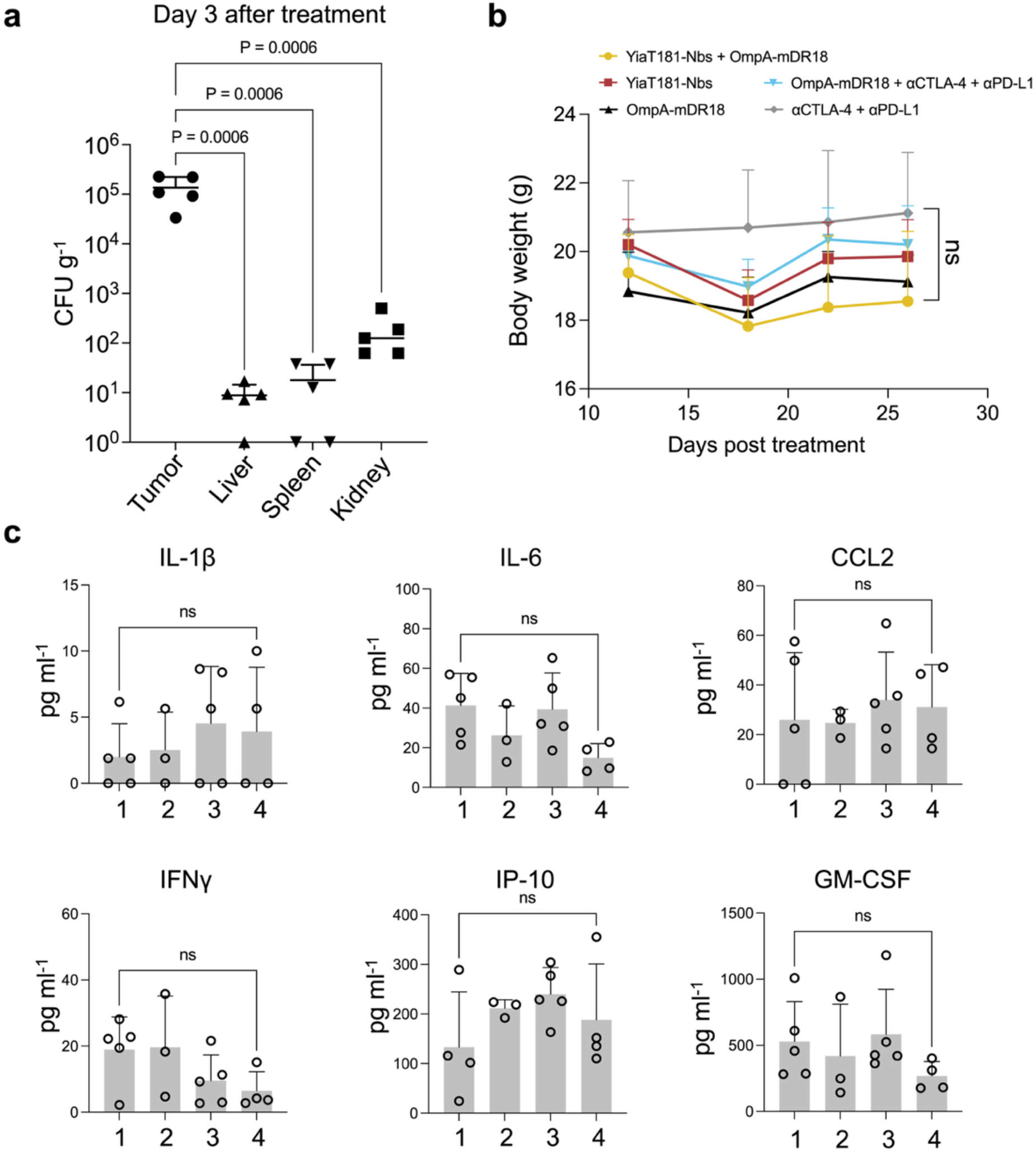
Biodistribution, safety assessment, and cytokine profiling following systemic administration of engineered bacteria in the B16F10 melanoma model. **a,** Biodistribution of engineered bacteria 3 days after intravenous administration. Bacterial burdens in tumors, liver, spleen, and kidney were quantified by colony-forming unit (CFU) enumeration and normalized to tissue weight. Each dot represents an individual mouse; horizontal lines indicate the mean. Statistical significance was determined by one-way ANOVA with Tukey’s multiple-comparisons test. **b,** Body weight measurements of B16F10 tumor-bearing mice following treatment with engineered bacterial therapies or antibody controls. Body weights were monitored throughout the study as an indicator of treatment tolerability. Data are presented as mean ± s.e.m. (n = 5 mice per group). Statistical significance was assessed using a mixed-effects model (REML). No significant differences were observed among treatment groups. **c,** Luminex-based quantification of cytokines and chemokines in plasma collected from B16F10 tumor-bearing mice 7 days after treatment (n = 3-5 mice per group). Concentrations of IL-1β, IL-6, CCL2, IFN-γ, IP-10/CXCL10, and GM-CSF were measured in mice treated with (1) YiaT181-Nbs + OmpA-mDR18, (2) YiaT181-Nbs, (3) OmpA-mDR18, or (4) the vehicle control, PBS. Data are presented as mean ± s.e.m. Other cytokines and chemokines are in **Extended Data Figures 5** and **6**. Each dot represents an individual mouse. Statistical significance was determined by one-way ANOVA with Tukey’s multiple-comparisons test.

### Bacterial membrane-anchored immunotherapy is associated with CD8⁺ T-cell enrichment and remodeling of the tumor immune microenvironment

Having established systemic antitumor activity, tumor-enriched bacterial localization, and limited systemic inflammatory toxicity, we next performed flow cytometric profiling of tumors and spleens 7 days after intravenous treatment in tumor-bearing mice. We analyzed both compartments because tumors represent the primary site where engineered bacteria accumulate and present membrane-anchored immunotherapeutic payloads, whereas spleens provide a complementary readout of systemic immune activation after intravenous bacterial administration (**Figure 5a, b**). Treatment groups included bacteria displaying (1) the combined YiaT181-Nbs and OmpA-mDR18, (2) YiaT181-Nbs alone, (3) OmpA-mDR18 alone, (4) commercial αCTLA-4 plus αPD-L1 antibodies, (5) commercial antibodies combined with OmpA-mDR18 bacteria, and (6) bacteria-only controls.

**Figure 5.**
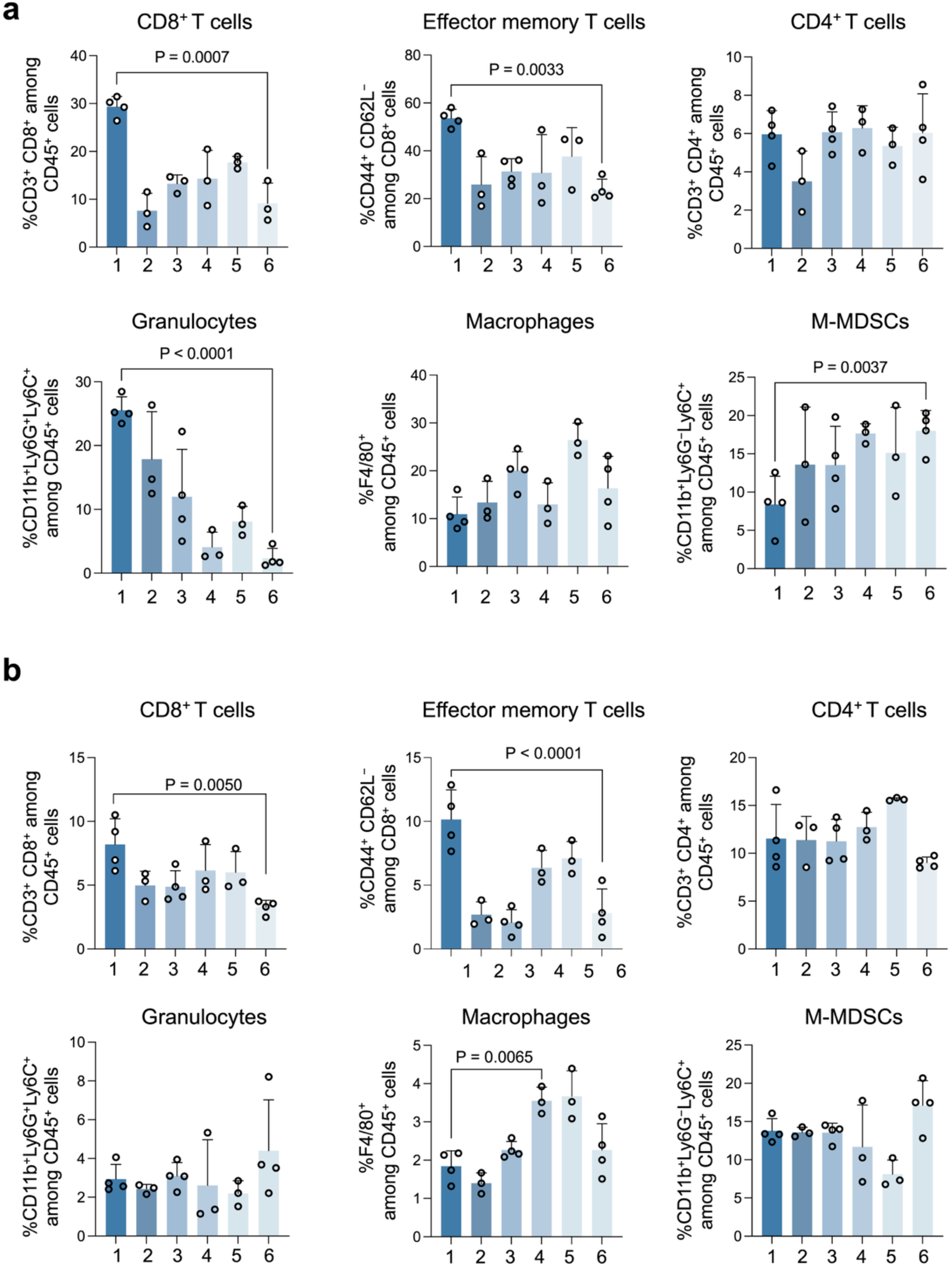
Bacterial membrane-anchored immunotherapy is associated with remodeling of the tumor immune microenvironment toward CD8⁺ T-cell immunity. Flow cytometric analysis of immune cell populations in MC38 tumors (**a**) and spleens (**b**) collected 7 days after intravenous administration of engineered bacteria. Quantification of CD8⁺ T cells, effector memory T cells (CD44⁺CD62L⁻ among CD8⁺ T cells), CD4⁺ T cells, granulocytes (CD11b⁺Ly6G⁺ among CD45⁺ cells), macrophages (F4/80⁺ among CD45⁺ cells), and monocytic myeloid-derived suppressor cells (M-MDSCs; CD11b⁺Ly6G⁻Ly6C⁺ among CD45⁺ cells). Data are presented as mean ± s.e.m. Each dot represents an individual mouse with n = 3 or 4 mice per treatment group. Statistical significance was determined by one-way ANOVA with multiple-comparison correction. P values are shown in the figure. Treatment groups were: (1) YiaT181-Nbs + OmpA-mDR18; (2) YiaT181-Nbs; (3) OmpA-mDR18; (4) commercial αCTLA-4 (clone 9D9) + αPD-L1 (clone 10F.9G2) antibodies; (5) OmpA-mDR18 + αCTLA-4 + αPD-L1; and (6) bacteria-only control.

Among the immune populations analyzed, the combined YiaT181-Nbs and OmpA-mDR18 treatment produced the most prominent increase in intratumoral CD8⁺ T cells compared with bacteria-only controls (**Figure 5a**). Effector-memory CD8⁺ T cells were also increased in this group, suggesting that membrane-anchored checkpoint blockade combined with bacterial surface-displayed mDR18 is associated with a more cytotoxic and memory-associated antitumor T-cell phenotype. In contrast, CD4⁺ T cells showed no major differences across treatment groups. The combination treatment also reshaped immunosuppressive and innate immune populations. Regulatory T cells were lower in the YiaT181-Nbs plus OmpA-mDR18 group than in mice receiving commercial antibody blockade combined with OmpA-mDR18, suggesting a more favorable effector-to-suppressor immune balance (**Extended Data Fig. 7**). Monocytic myeloid-derived suppressor cells (M-MDSCs) were also reduced in the combined bacterial treatment group compared with antibody-based treatment groups and bacteria-only controls. Granulocytes were enriched in the combined bacterial treatment group, whereas macrophages, monocytes, and NK cells showed less pronounced changes across groups.

Spleen immune profiling revealed similar immune changes after systemic treatment (**Figure 5b**). The increased CD8⁺ and effector-memory CD8⁺ T-cell populations in spleens suggest that intravenous bacterial therapy can also engage systemic adaptive immunity, while the absence of broad inflammatory immune expansion is consistent with the limited systemic cytokine induction observed in plasma. Together, the tumor and spleen immune-profiling data support the conclusion that systemically delivered YiaT181-Nbs plus OmpA-mDR18 is associated with coordinated remodeling of local tumor immunity and systemic adaptive immune activation.

## Discussion

In our previous work, non-pathogenic, tumor-homing *E. coli* were engineered to display decoy-resistant IL-18 on the bacterial outer membrane, enabling localized cytokine stimulation, CD8⁺ T cell- and NK cell-dependent antitumor immunity, and enhanced CAR-NK cell activity in solid tumors ^27^. In a separate study, surface display was applied to a distinct disease mechanism, where engineered bacteria displaying the colibactin resistance protein ClbS neutralized genotoxic colibactin in the gut and reduced *pks*⁺ *E. coli*-driven intestinal tumorigenesis ^45^. Together, these studies established bacterial surface display as a versatile strategy for positioning therapeutic proteins at disease-relevant interfaces. The present work extends this concept to immune checkpoint blockade by engineering tumor-homing *E. coli* strains that each express checkpoint-blocking nanobodies targeting CTLA-4 or PD-L1. Unlike secretion- or lysis-based bacterial delivery systems, this approach anchors checkpoint blockers directly on the bacterial outer membrane. This design is intended to exploit bacterial enrichment in tumors, increase local avidity, spatially restrict checkpoint blockade, and potentially reduce systemic exposure. The systemic delivery results support this rationale: intravenously administered bacteria preferentially accumulated in tumors over liver, spleen, and kidney, and the combined YiaT181-Nbs plus OmpA-mDR18 therapy did not induce broad systemic cytokine elevation. These findings are consistent with the hypothesis that bacterial tropism and membrane anchoring can concentrate biologic activity within tumors while limiting systemic inflammatory toxicity.

A central finding in our study is that scaffold choice strongly influences therapeutic function. Although Neae supported high display levels, YiaT181 resulted in stronger PD-L1 binding and superior antitumor activity, whereas OmpA was less effective for checkpoint nanobody display. This contrasts with our prior DR-18 work, where OmpA was an effective scaffold for cytokine display ^27^. A similar observation was made in the ClbS study, in which OmpA-mediated display enabled higher antitoxin activity ^45^. These comparisons highlight an important design principle: optimal scaffold selection is payload dependent. Display abundance alone is insufficient; protein orientation, folding, target accessibility, and receptor geometry are also likely to determine biological activity. Our use of MC38 and B16F10 further allowed us to evaluate bacterial surface-displayed PD-L1 blockade across tumor contexts in which the dominant source of immunosuppressive PD-L1 may differ. In MC38, prior genetic studies showed that tumor-cell PD-L1 can directly suppress CD8⁺ T-cell cytotoxicity and confer a selective advantage *in vivo*, supporting the relevance of targeting tumor-associated PD-L1. In contrast, B16 melanoma models highlight the importance of PD-L1 expressed by non-tumor host cells ^30^, suggesting that a tumor-localized bacterial display platform may be advantageous because it can position checkpoint-blocking nanobodies near both malignant cells and PD-L1⁺ host immune or stromal cells within the tumor microenvironment. These observations support the broader rationale that membrane-anchored bacterial checkpoint blockade may intercept distinct immune-suppressive mechanisms mediated by PD-L1 at tumor sites. Future studies using cell-type-specific PD-L1 analysis or genetic models will help define which PD-L1-expressing cellular compartments are most directly engaged by this platform in each tumor model.

A major advance of the current work is the therapeutic integration of different surface-displayed immunotherapy modules through co-administration of single-payload bacterial strains. By combining YiaT181-displayed checkpoint nanobody strains with a separate OmpA-displayed mDR18 strain, we show that bacterial surface display can support modular assembly of complementary immune functions on living therapeutics. The combined bacterial treatment suppressed tumor growth more effectively than either component alone and outperformed benchmark antibody-based checkpoint blockade under the tested conditions. It also promoted tumor rejection and rechallenge resistance, while immune profiling showed increased CD8⁺ and effector-memory T-cell responses together with reduced suppressive myeloid populations. However, several limitations remain to be addressed. We reason that the performance of different outer membrane proteins may vary across nanobody sequences. Furthermore, future studies should examine the relative contributions of CTLA-4 blockade, PD-L1 blockade, bacterial innate immune activation, and DR18 signaling, which will require further mechanistic dissection. For example, depletion or genetic perturbation studies will help determine whether CD8⁺ T cells, NK cells, or other immune populations are required for therapeutic efficacy. Future work may also address genetic stability via genomic integration rather than relying on plasmid maintenance, bacterial persistence and clearance, and safety control for systemic administration. Overall, this work advances bacterial surface display as a modular strategy for spatially confined biologic therapy at tumor and host-microbe interfaces.

## Acknowledgments

This work was supported by Swim Across America (SAA) Young Investigator Award (JL), the NIH Office of the Director (1DP2GM154019-01) (JL), and the National Cancer Institute (R01CA299949) (JL). We want to thank the ULAM Pathology Core and the Flow Cytometry Core at the University of Michigan. We would like to express our gratitude to the other members of the Li lab and Indrani Talukder from the Shah lab for providing essential resources and helpful suggestions.

## Author contributions

J.L. and Z.W. conceptualized the study. J.L. and Z.W. wrote the manuscript. J.L., Z.W., C.F., X.Z. designed the experiments, analysis, and data interpretation. Z.W., C.F., X.Z. performed the experiments. S.Y., J.J.M., R.R., and Y.S. provided guidance during optimization and assisted with experimental procedures and analysis. All the authors contributed to data interpretation and the final version of the manuscript.

## Materials and methods

### Cell culture

MC38 and Yummer1.7 were obtained from Dr. Darrell Irvine’s laboratory at the Koch Institute for Integrative Cancer Research (Cambridge, MA, USA). The DC 2.4 cell line was obtained from Kenneth Rock (University of Massachusetts, Worcester, MA). B16F10 (CRL-6475) was purchased from American Type Culture Collection (ATCC, Manassas, VA, USA). MC38, B16F10, and DC2.4 cells were maintained in Dulbecco’s modified Eagle’s medium (DMEM; Corning, NY, USA) supplemented with 10% fetal bovine serum (FBS; Corning) and 100 U ml⁻¹ penicillin-streptomycin (Corning). Yummer1.7 cells were maintained in DMEM supplemented with 10% FBS, 100 U ml⁻¹ penicillin-streptomycin, and 1× non-essential amino acids (NEAA; Gibco, Grand Island, NY, USA). All cell lines were cultured at 37 °C in a humidified incubator containing 5% CO₂. Cells between passages 2 and 10 were used for all experiments. All cell lines were routinely tested and confirmed negative for mycoplasma contamination.

### Escherichia coli surface display

For bacterial surface display of mDR18 or checkpoint nanobodies targeting CTLA-4 (PDB: 5E03) and PD-L1 (PDB: 5DXW), we used the following plasmid backbones: pDSG323 (Addgene #115594) ^37^ with C-terminal fusion scaffold protein Neae, pDS861 (a generous gift from Quintara Biosciences) with C-terminal fusion scaffold protein OmpA or YiaT181, pHEA (Addgene #168297) ^38^, which utilizes EhaA, an autotransporter protein of enterohemorrhagic *Escherichia coli* O157:H7 that contributes to adhesion and biofilm formation, and pAIDA1 (Addgene #79180) ^39^, which utilizes the adhesin protein involved in diffuse adherence of enteropathogenic *E. coli*. For C-terminal display scaffolds, nanobodies were fused downstream of the scaffold through three GGGGS linkers. For N-terminal autotransporter display scaffolds, nanobodies were fused upstream of the transporter domain through the corresponding linker architecture. DNA sequences for the encoding nanobodies, GGGGS linker, and DYKDDDDK-tag (FLAG-tag) were inserted between *SpeI* and *PstI* sites for pDSG323, between *NotI* and *BamHI* sites for pDS861, between *NotI* and *EcoRI* sites for pHEA, and between *KpnI* and *XbaI* sites for pAIDA1 by NEBuilder® HiFi DNA Assembly Master Mix (NEB, catalog#: M5520AVIAL). For the optimization of the surface display of OmpA-Nb, the high-copy plasmid pDS861-dsRed (from Quintara Biosciences) was digested with *XbaI* and *NotI*. The *E. coli* rhaSR-PrhaBAD inducible promoter system, including RhaR/RhaS regulatory proteins, the rhamnose-responsive PrhaBAD promoter, and a ribosome-binding site, was assembled into the digested pDS861-dsRed backbone using NEBuilder HiFi DNA Assembly. The DNA fragments for nanobodies with an N-terminal Myc tag and a C-terminal FLAG-tag were synthesized by Integrated DNA Technologies (IDT, Coralville, IA), or 3 copies of GGGGS linkers were introduced by primers. All plasmids were validated by Whole Plasmid Sequencing before the next steps.

For bacterial induction, *E. coli* K-12 DH5α with the corresponding plasmid was inoculated into fresh Luria-Bertani (LB) medium containing 50 μg/ml kanamycin or 25 μg/ml chloramphenicol. After overnight culture in a shaker (37 °C, 250 rpm), bacterial suspensions were diluted 10-fold in the fresh LB with 50 μg/ml kanamycin and 10 mM L-Rhamnose for derivatives of pDS861, in the fresh LB with 50 μg/ml kanamycin and 100 ng/ml anhydrotetracycline (aTc) for modified plasmids based on pDSG323, and in fresh LB medium supplemented with 25 μg/ml chloramphenicol and 1 mM isopropyl β-D-1-thiogalactopyranoside (IPTG) for derivatives of pHEA and pAIDA1. After 24 h of induction in a shaker (25 °C, 250 rpm), bacteria were collected for the next step. For bacterial surface display verification, 20 μl bacterial suspension was collected, washed once with phosphate-buffered saline (PBS), and incubated with anti-DYKDDDDK Tag antibody (BioLegend, San Diego, CA, USA; catalog no. 637315) for FLAG-tagged constructs or MonoRab™ Rabbit Anti-Camelid VHH Cocktail [iFluor 647] (GenScript, Piscataway, NJ, USA; catalog no. A02019) for nanobody-displaying constructs in FACS buffer (2% FBS, 0.1% w/v sodium azide, 2 mM Ethylenediaminetetraacetic acid dissolved in DPBS) for 20 min at room temperature, washed twice and then suspended in PBS for flow cytometry (Attune NxT Flow Cytometer, Thermo Fisher Scientific, Waltham, MA, USA). A red laser, along with a 670-nm/14-nm filter, was used.

### Bacteria-host cell binding assay

After induction, bacteria expressing each scaffold displaying nanobodies were harvested by centrifugation and washed twice with PBS. Bacteria were stained with 5 µM CFSE (Tonbo Biosciences, San Diego, CA, USA; Cat# 13-0850-U500) at 37 °C for 15 minutes, washed twice with PBS, and resuspended in cell culture medium. Mouse DC2.4 dendritic cells were seeded at 4 × 10⁴ cells per well in 96-well cell culture plates the day before coculture. The next day, induced bacterial cultures were pelleted, washed twice with sterile PBS, and resuspended at the indicated concentration. Bacterial density was estimated by OD600 using a formula OD600 = 1.0 = 8 × 10⁸ CFU/ml. Prior to coculture, DC2.4 cells were washed twice with DPBS. CFSE-labeled bacteria were incubated with DC2.4 cells at the indicated bacteria-to-cell ratio (typically MOI = 100:1). To assess bacterial binding while minimizing bacterial internalization, cocultures were incubated at 4 °C for 30 min. Following incubation, cells were washed extensively with PBS to remove unbound bacteria and analyzed by flow cytometry using the Attune NxT Flow Cytometer. CFSE fluorescence was detected using the 488-nm blue laser and a 530/30-nm emission filter. Bacterial association with DC2.4 cells was quantified based on the percentage of CFSE-positive cells and the mean fluorescence intensity (MFI) of the CFSE signal. Relative binding was determined by normalizing the signal of nanobody-displaying strains to that of the corresponding scaffold-only control strain (OmpA, YiaT181, or Neae). B16F10 melanoma cells were used in parallel to validate bacterial binding under the same experimental conditions.

### Recombinant CTLA-4 binding assay

Engineered *E. coli* expressing anti-mouse CTLA-4 nanobodies were grown overnight in LB medium at 37 °C with shaking, then diluted 1:10 into fresh LB medium containing the appropriate inducer. Cultures were induced for 24 h at 25 °C, harvested by centrifugation, washed twice with PBS, and resuspended in PBS supplemented with 1% bovine serum albumin (BSA). Approximately 1 × 10⁸ bacteria were incubated with biotinylated recombinant mouse CTLA-4 protein (Sino Biological, Beijing, China; Cat. No. 50503-M49H-B) at the indicated concentration (1 μg/ml) for 20 min at room temperature. Following incubation, bacteria were washed twice with PBS and stained with FITC-conjugated streptavidin (BioLegend, San Diego, CA, USA; Cat. No. 405202) at a 1:200 dilution for 20 min at room temperature in the dark. Bacteria were subsequently washed twice with PBS and analyzed by flow cytometry (Invitrogen Attune). Blue laser, along with a 530/30 nm filter, was used to detect FITC. Binding of recombinant CTLA-4 protein to surface-displayed αCTLA-4 nanobodies was quantified based on the MFI of the FITC signal. Scaffold-only strains were included as negative controls in all experiments. Relative binding was determined by normalizing the fluorescence signal of nanobody-displaying strains to that of the corresponding scaffold-only control strain.

### Animal experiments

Female C57BL/6J mice (6–8 weeks old; JAX stock no. 000664) were purchased from The Jackson Laboratory (Bar Harbor, ME, USA) and maintained in the Unit for Laboratory Animal Medicine at the University of Michigan. Mice were housed under specific pathogen-free (SPF) conditions in a temperature- and humidity-controlled facility with a 12-h light/12-h dark cycle and provided ad libitum access to standard chow and water. All animal experiments were conducted in accordance with federal, state, and institutional guidelines and were approved by the Institutional Animal Care and Use Committee (IACUC) of the University of Michigan. All procedures were performed in compliance with the Association for Assessment and Accreditation of Laboratory Animal Care International (AAALAC) guidelines.

Female C57BL/6J mice were inoculated subcutaneously in the flank with 0.5 × 10^6^ MC38 or Yummer1.7 cells in 50 μL sterile PBS, or 0.5 × 10^6^ B16F10 cells in 50 μL sterile PBS, as indicated for each experiment. Tumor dimensions were measured every 3 days using digital calipers, and tumor volume was calculated as length × width² / 2, where length represents the longest tumor dimension, and width represents the perpendicular dimension. Mice were randomized to treatment groups when tumors reached the indicated size range.

For scaffold screening studies, mice bearing MC38 tumors (50–100 mm³) received intratumoral injections of sterile PBS or engineered bacteria (20 μL) twice weekly for a total of 2 doses. For intratumoral efficacy studies, mice bearing MC38 or Yummer1.7 tumors were treated when tumors reached 50–100 mm³. Mice received intratumoral injections of PBS or engineered bacteria (2.5 × 10^8^ CFU per strain) in 20 μL sterile PBS twice weekly for a total of two doses. Scaffold-only groups received 5 × 10⁸ CFU per dose. Nbs-treated groups received a total of 5 × 10⁸ CFU per dose, consisting of 2.5 × 10⁸ CFU αCTLA-4 Nb-displaying bacteria and 2.5 × 10⁸ CFU αPD-L1 Nb-displaying bacteria. Combination therapy groups received Nbs together with 2.5 × 10⁸ CFU OmpA-mDR18, for a total dose of 7.5 × 10⁸ CFU in the treatment.

For systemic delivery studies, mice bearing Yummer1.7 or B16F10 tumors (120–200 mm³) were randomized to receive PBS, engineered bacteria (3.3×10^8^ CFU per strain), anti-mouse PD-L1 antibody (clone 10F.9G2, Bio X Cell, Lebanon, NH, USA; BE0101), anti-mouse CTLA-4 antibody (clone 9D9, Bio X Cell, Lebanon, NH, USA; BE0164), or a combination of engineered bacteria and checkpoint-blocking antibodies. Specifically, YiaT181-Nbs-treated mice received a total of 6.6 × 10⁸ CFU per dose, consisting of 3.3 × 10⁸ CFU of each nanobody-displaying strain. OmpA-mDR18-treated mice received 3.3 × 10⁸ CFU per dose. The bacterial combination group received 3.3 × 10⁸ CFU of each strain, corresponding to a total dose of approximately 1 × 10⁹ CFU. Antibody-treated groups received 200 μg anti-CTLA-4 and 100 μg anti-PD-L1 per mouse. Engineered bacteria were administered intravenously in 100 μL sterile PBS, whereas antibodies were administered intraperitoneally in 100 μL sterile PBS. Treatments were administered twice weekly for a total of three doses. In all animal studies, mice were monitored daily following intravenous bacterial administration for signs of toxicity, including body weight loss, lethargy, impaired mobility, and other indicators of morbidity. Mice were euthanized upon reaching institutional humane endpoint criteria, including tumor volume exceeding 1,500 mm³, or >20% body weight loss.

### Biodistribution and bacterial enumeration

To assess bacterial biodistribution, mice were euthanized at the indicated time points following bacterial administration. Tumors, livers, spleens, and kidneys were aseptically harvested, weighed, and transferred to sterile tubes containing cold PBS. Tissues were homogenized using a bead homogenizer and serially diluted in PBS. Aliquots (2 μl) of each dilution were plated on LB agar supplemented with the appropriate antibiotics to selectively recover engineered bacteria. Plates were incubated at 37 °C overnight, and colony-forming units (CFUs) were enumerated the following day. Bacterial burden was calculated as CFU per gram of tissue using the following formula: CFU/g = (number of colonies × dilution factor × homogenization volume) / tissue weight. For samples with no detectable colonies, values were assigned to the limit of detection for statistical analysis. The limit of detection was determined by the minimum number of colonies detectable from the plated homogenate volume and is indicated in the corresponding figure legends. Biodistribution data are presented as CFU/g tissue.

### Luminex analysis

Blood and tumor samples were collected from tumor-bearing mice 7 days after intravenous administration of engineered bacteria. Whole blood was collected by cardiac puncture into tubes containing EDTA anticoagulant and centrifuged to isolate plasma. Plasma samples were aliquoted and stored at −80 °C until analysis. For tumor lysate preparation, approximately 20 mg of freshly excised tumor tissue was lysed in T-PER Tissue Protein Extraction Reagent (Thermo Fisher Scientific) supplemented with protease and phosphatase blockers. Tumor samples were incubated on a rotator at 4 °C for 30 min and subsequently centrifuged at maximum speed at 4 °C to remove insoluble debris. The clarified supernatants were collected and stored at −80 °C until analysis. Total protein concentrations were determined using the DC Protein Assay (Bio-Rad Laboratories), and samples were normalized based on total protein concentration prior to cytokine analysis. Cytokine measurements from tumor lysates were normalized to total protein content. Tumor lysate and plasma samples were submitted to Eve Technologies (Calgary, Alberta, Canada) for cytokine analysis using the Mouse Cytokine/Chemokine 32-Plex Discovery Assay. Multiplex quantification was performed using Luminex xMAP technology (Luminex 200, DiaSorin) according to the manufacturer’s instructions (MILLIPLEX Mouse Cytokine/Chemokine Magnetic Bead Panel, MilliporeSigma). Cytokine concentrations were calculated from standard curves and reported as pg/ml.

### Immune cell profiling in mice

Tumor and spleen tissues were collected from tumor-bearing mice 7 days after bacterial treatment and processed into single-cell suspensions for immune profiling. Tumors were minced into small fragments and dissociated using a gentleMACS tissue dissociator (Miltenyi Biotec), followed by filtration through a 70 μm cell strainer and washing with 1% BSA in PBS. Spleens were mechanically dissociated by gently mashing through a 70 μm cell strainer using a syringe plunger. Following centrifugation, red blood cells were lysed using ACK lysis buffer (Gibco, A10492), and the resulting cell suspensions were filtered through a 70 μm strainer and washed with FACS buffer. For flow cytometric staining, single-cell suspensions were first stained with a viability dye (LIVE/DEAD Fixable Near-IR, Invitrogen, L10119) and incubated with Fc blocking antibody (anti-mouse CD16/CD32, Fisher Scientific, 50-112-9520). Cells were subsequently stained with surface antibody cocktails for 30 min at room temperature. For intracellular staining, cells were fixed and permeabilized using the Fixation/Permeabilization Kit (BD Biosciences, 554714) according to the manufacturer’s instructions, then incubated with intracellular antibody cocktails for 30 min at room temperature. Cells were then washed twice and resuspended in FACS buffer for acquisition. Flow cytometry data were acquired on a Cytek Aurora spectral flow cytometer (Cytek Biosciences, Fremont, CA, USA) at the University of Michigan Flow Cytometry Core using SpectroFlo software. Antibodies were used at a dilution of 1:200 for surface staining and 1:100 for intracellular staining unless otherwise indicated. The antibodies used for immune profiling are listed in **Table S4**, and the gating strategy is shown in **Extended Data Fig. 4.** Data were analyzed using FlowJo v11.1.1 software (BD Biosciences, Ashland, OR, USA).

### Statistical Analysis

Statistical analyses were performed using GraphPad Prism for macOS (version 11.0.2, GraphPad Software, San Diego, CA, USA). Specific statistical tests are indicated in the corresponding figure legends. All tests were two-sided unless otherwise specified. P values < 0.05 were considered statistically significant. Data are presented as mean ± s.e.m. unless otherwise indicated.

## Data Availability

Raw data, source data, plasmid maps/sequences, and flow cytometry gating files/source data are available upon request.

## Extended Data Figures

**Extended Data Figure 1.**
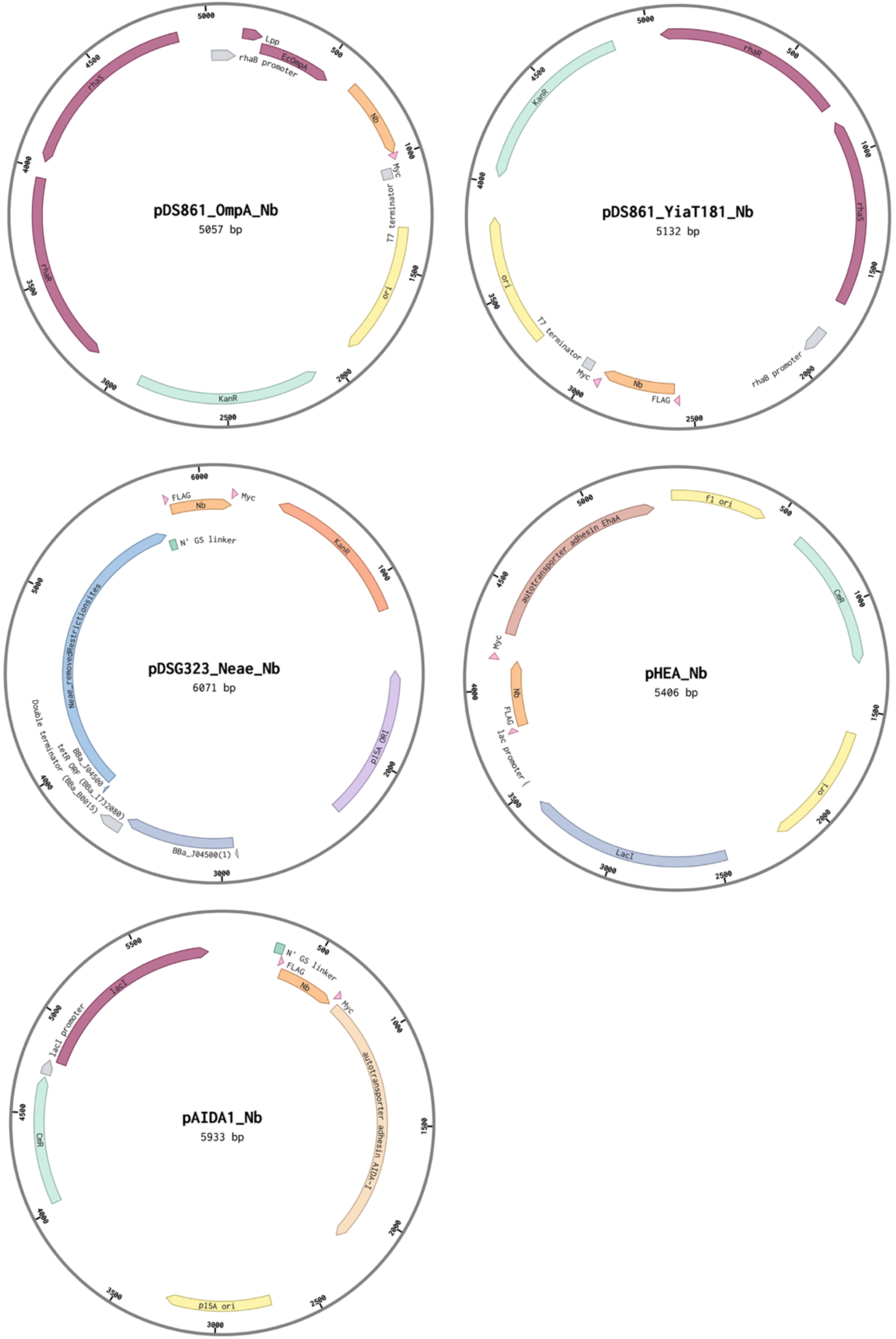
Plasmid maps used for displaying Nbs by OmpA, YiaT181, Neae, C-EhaA and AIDA. Full plasmid sequences will be provided upon request.

**Extended Data Figure 2.**
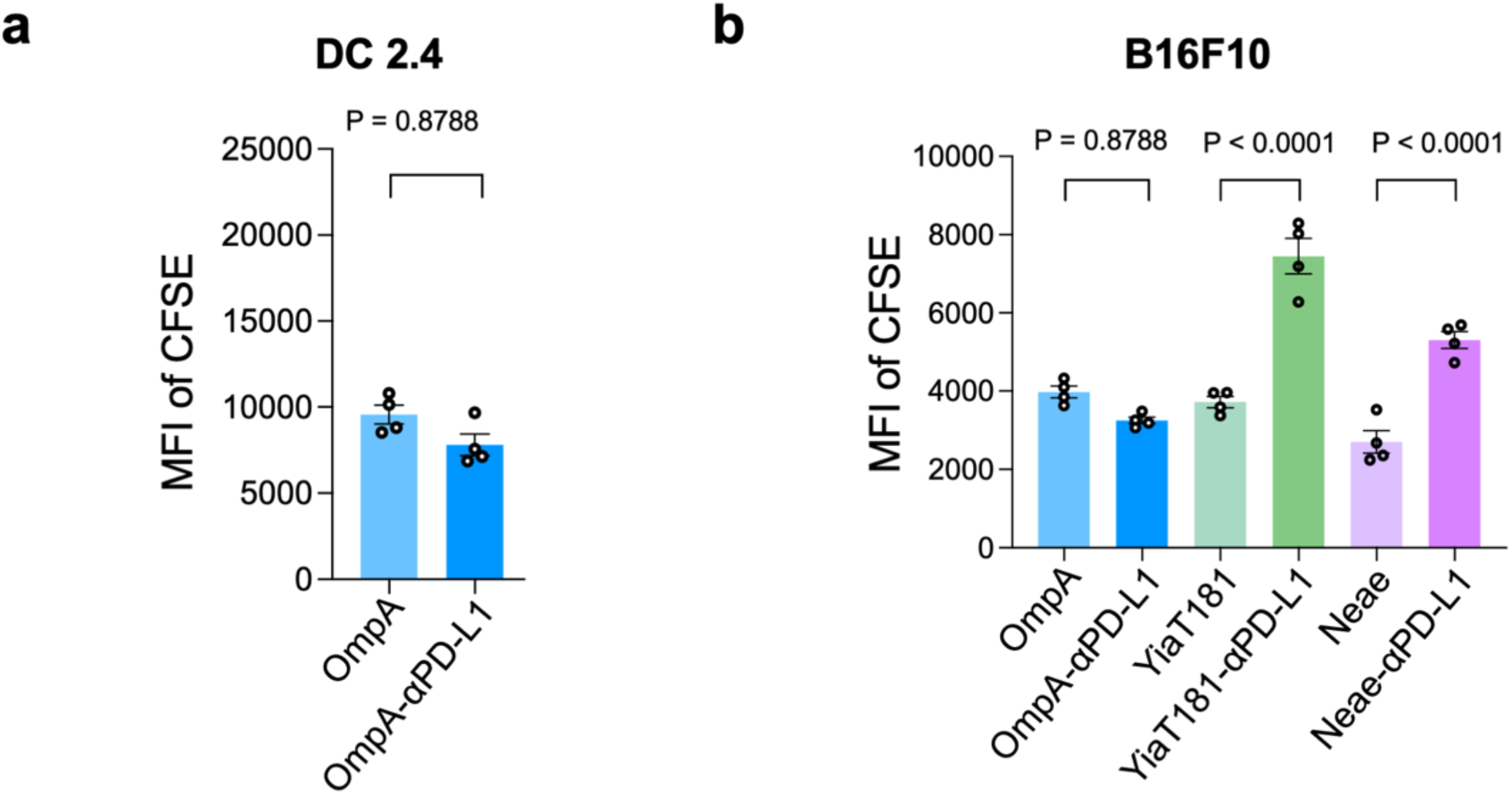
Evaluation of bacterial surface display scaffolds for PD-L1 nanobody-mediated binding to target cells. **a,** Quantification of binding between engineered *E. coli* displaying αPD-L1 nanobodies through the OmpA scaffold and mouse DC 2.4 dendritic cells. CFSE-labeled bacteria were incubated with DC2.4 cells, and bacterial association was quantified by flow cytometry as the mean fluorescence intensity (MFI) of CFSE-positive events. Bars represent mean ± s.e.m. Each dot represents an individual technical repeat (n = 4) from two independent experiments. Statistical significance was determined using an unpaired two-tailed t-test. **b,** Comparison of PD-L1 nanobody-mediated binding efficiency among different bacterial surface display scaffolds. CFSE-labeled engineered *E. coli* expressing αPD-L1 nanobodies through OmpA, YiaT181, or Neae were incubated with B16F10 melanoma cells and analyzed by flow cytometry. Binding was quantified as the MFI of CFSE-positive events. Bars represent mean ± s.e.m. Each dot represents an individual technical repeat (n = 4) from two independent experiments. Statistical significance was determined by one-way ANOVA with multiple-comparison correction. P values are indicated in the figure.

**Extended Data Figure 3.**
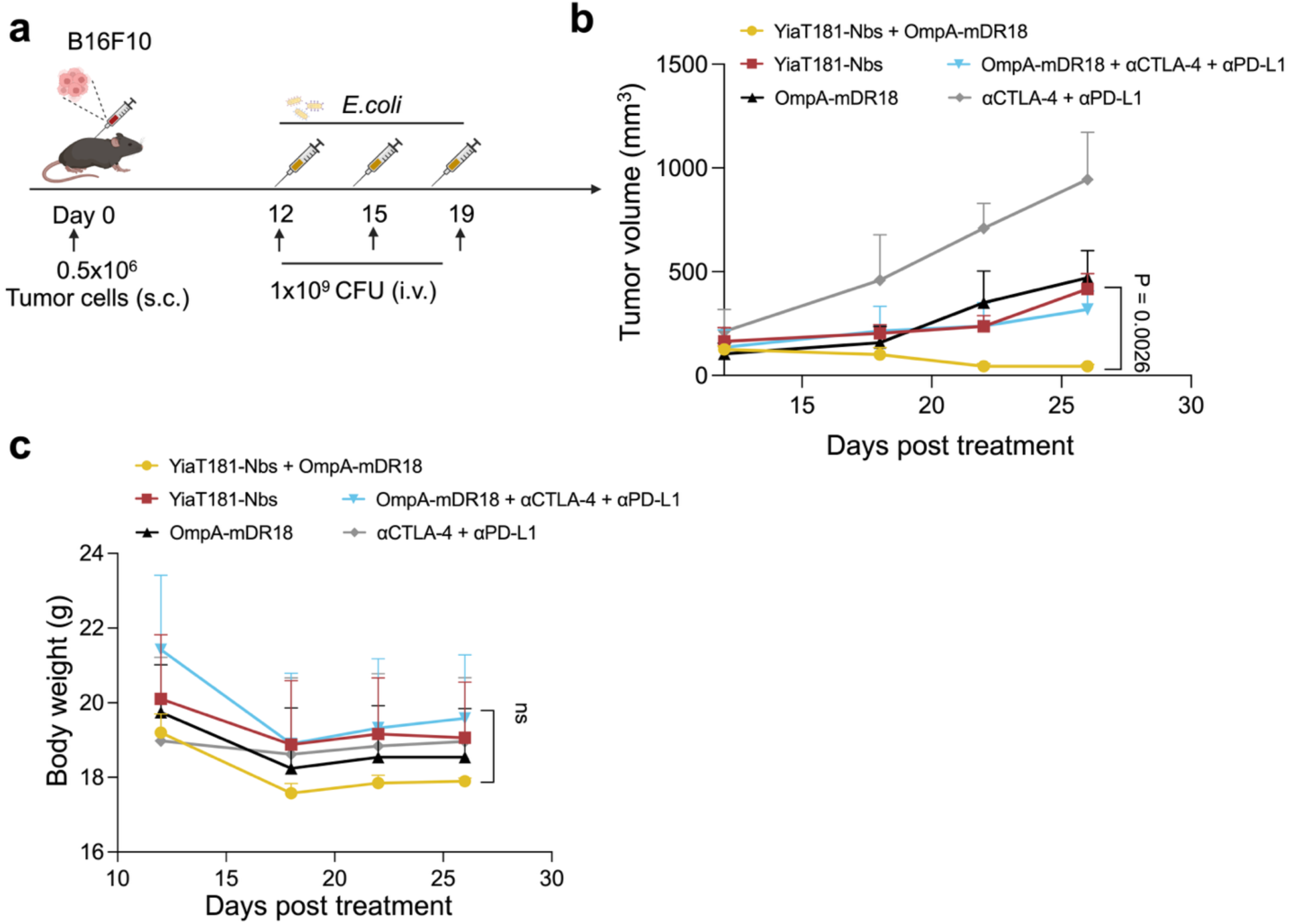
Systemic administration of a combination of engineered bacteria, each displaying a separate checkpoint nanobody or mDR18, suppresses tumor growth in the B16F10 melanoma model. **a,** Schematic illustration of the treatment regimen in B16F10 tumor-bearing mice. Mice were subcutaneously inoculated with 5 × 10⁵ B16F10 melanoma cells on day 0. Engineered *E. coli* were administered intravenously at a dose of 10⁹ CFU on days 12, 15, and 19 after tumor implantation. **b,** Tumor growth curves of B16F10 tumor-bearing mice treated with engineered bacteria displaying YiaT181-Nbs, OmpA-mDR18, or the combination of both. Additional control groups received systemic αCTLA-4 and αPD-L1 antibodies alone or in combination with OmpA-mDR18 bacteria. Tumor volumes were measured at the indicated time points and are presented as mean ± s.e.m. (n = 5 mice per group). Statistical significance was determined by two-way ANOVA followed by Tukey’s multiple-comparisons test. The indicated P value compares the combination treatment group with the relevant control groups. **c,** Body weight measurements of B16F10 tumor-bearing mice following treatment with engineered bacterial therapies or antibody controls. Body weights were monitored throughout the study as an indicator of treatment tolerability. Data are presented as mean ± s.e.m. (n = 5 mice per group). Statistical significance was assessed using a mixed-effects model (REML). No significant differences were observed among treatment groups.

**Extended Data Figure 4.**
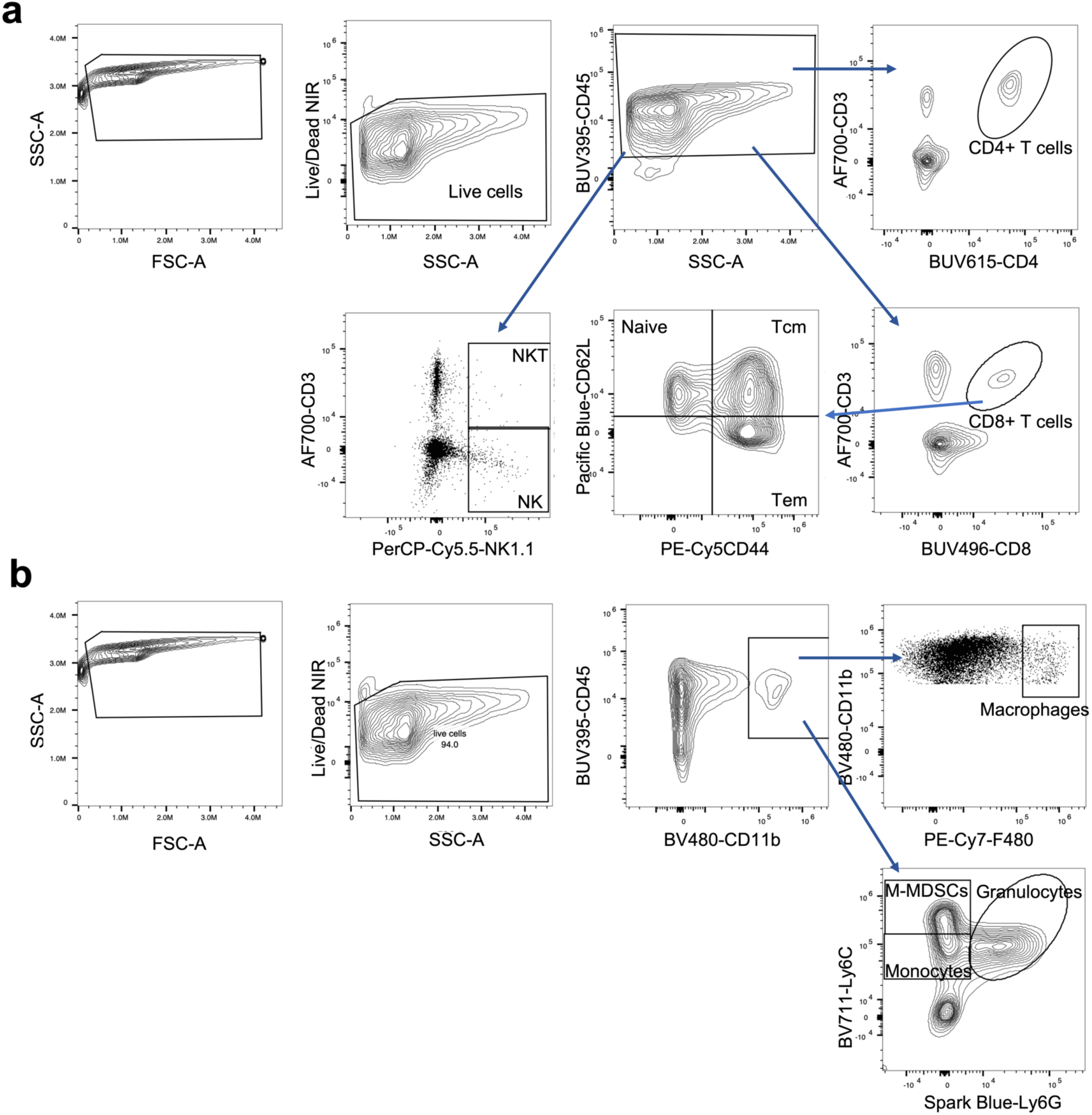
Representative gating strategy of major immune cell populations in the tumor and spleen of C57BL/6J mice. **a,** Gating strategy for lymphoid populations. Live CD45⁺ leukocytes were identified and subsequently gated to define CD4⁺ T cells, CD8⁺ T cells, and T-cell memory subsets, including naïve (CD44⁻CD62L⁺), central memory (Tcm; CD44⁺CD62L⁺), and effector memory (Tem; CD44⁺CD62L⁻) T cells. **b,** Gating strategy for myeloid populations. Live CD45⁺CD11b⁺ cells were further analyzed to identify macrophages (F4/80⁺), monocytes (Ly6C⁺Ly6G⁻), monocytic myeloid-derived suppressor cells (M-MDSCs; Ly6C^high^Ly6G⁻), and granulocytes (Ly6G⁺Ly6C^low^).

**Extended Data Figure 5.**
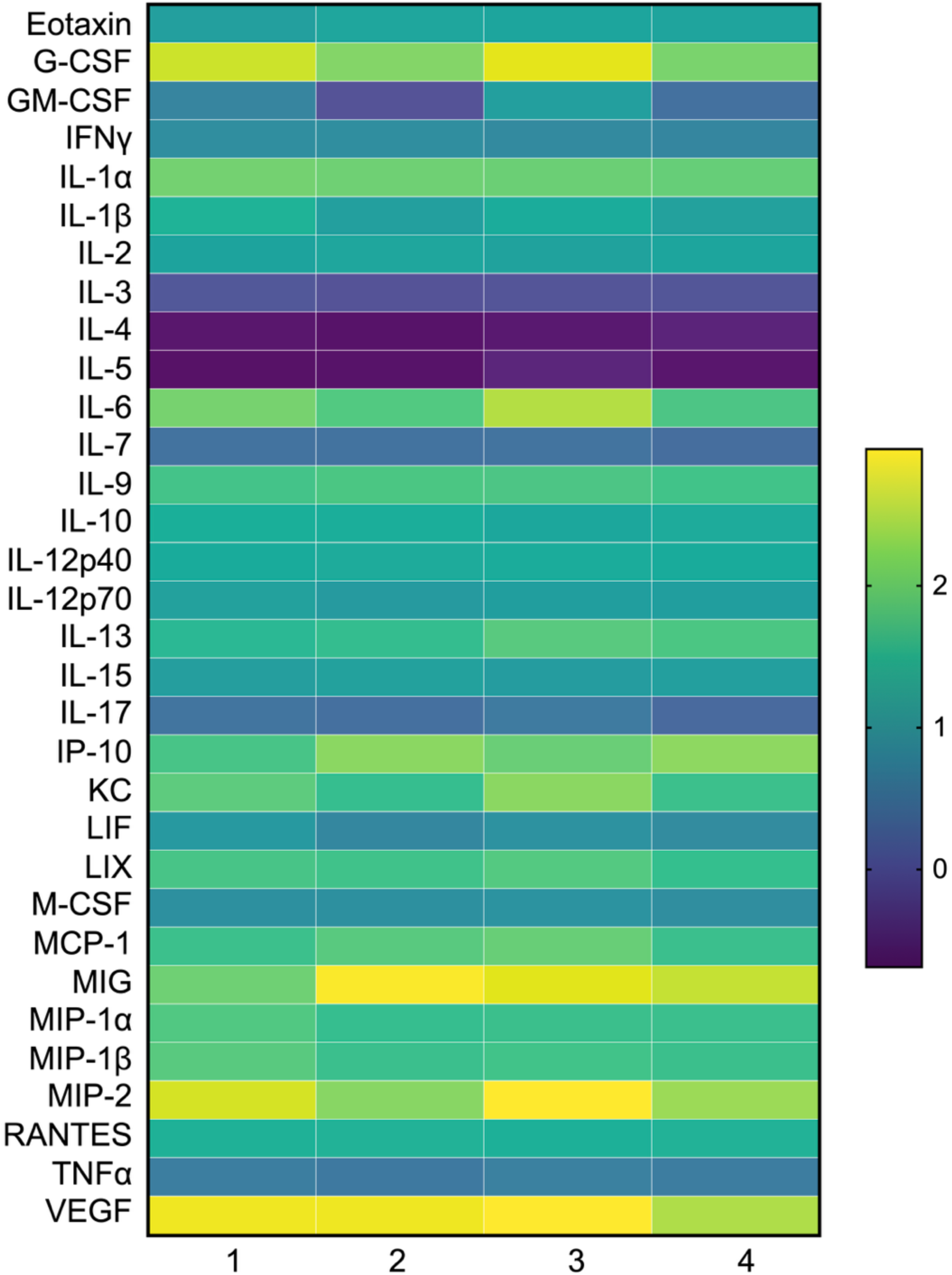
Cytokine/chemokine profiling of B16F10 tumor homogenates following systemic treatment with engineered bacteria. Heatmap showing median cytokine and chemokine concentrations in tumor homogenates collected from B16F10 melanoma-bearing mice 7 days after intravenous administration of engineered bacteria (n = 3-5 mice per group). Cytokine levels were quantified using a 32-plex Luminex assay and are presented as log10-transformed median concentrations (pg mL⁻¹). Treatment groups were: (1) YiaT181-Nbs + OmpA-mDR18, (2) YiaT181-Nbs, (3) OmpA-mDR18, and (4) PBS.

**Extended Data Figure 6.**
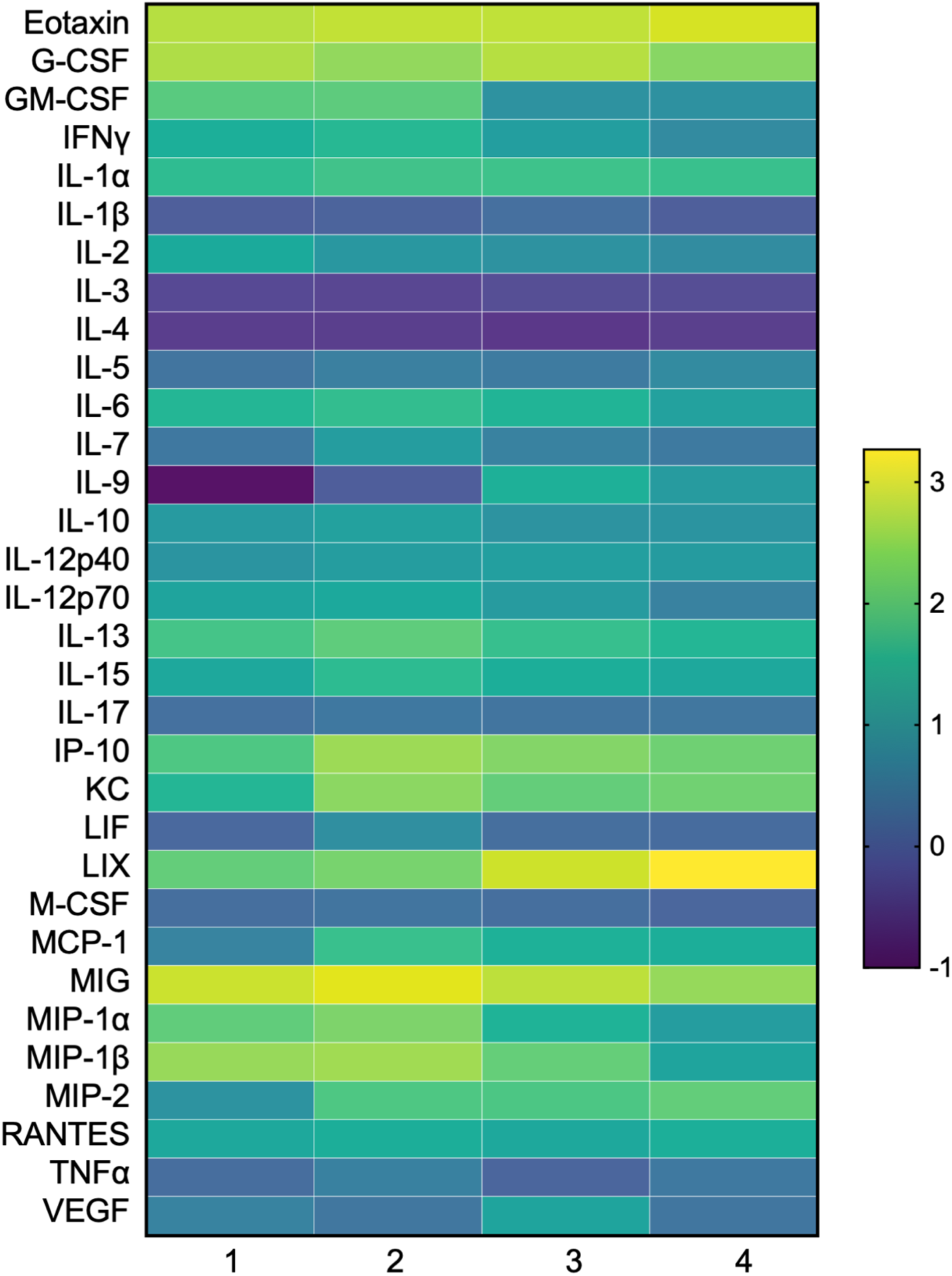
Cytokine/chemokine profiling in mouse plasma following systemic treatment with engineered bacteria in the B16F10 melanoma model. Heatmap showing median cytokine and chemokine concentrations in plasma collected from B16F10 melanoma-bearing mice 7 days after intravenous administration of engineered bacteria (n = 3-5 mice per group). Cytokine levels were quantified using a 32-plex Luminex assay and are presented as log10-transformed median concentrations (pg mL⁻¹). Treatment groups were: (1) YiaT181-Nbs + OmpA-mDR18, (2) YiaT181-Nbs, (3) OmpA-mDR18, and (4) PBS.

**Extended Data Figure 7.**
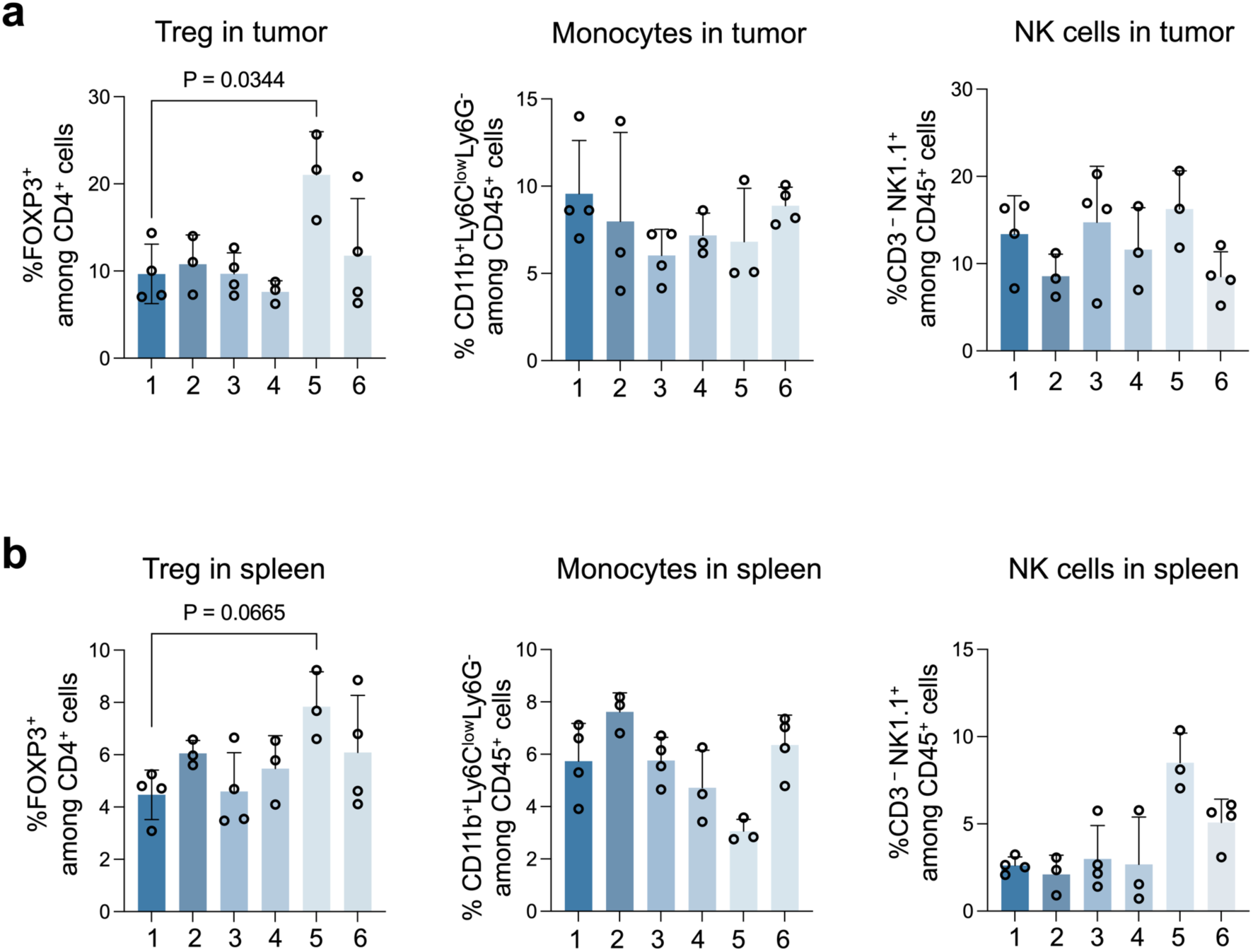
Quantification of regulatory T cells, monocytes, and natural killer cells in tumors and spleens following bacterial therapy. Frequencies of regulatory T cells (Tregs; FOXP3⁺ cells among CD4⁺ T cells), monocytes (CD11b⁺Ly6C^low^Ly6G⁻ cells among CD45⁺ cells), and natural killer (NK) cells (CD3⁻NK1.1⁺ cells among CD45⁺ cells) in **a,** tumors or **b,** spleens from MC38 tumor-bearing mice 7 days after systemic delivery of the indicated bacterial therapies (n = 3-4 mice per group). Data are presented as mean ± s.e.m. Each dot represents an individual mouse. Statistical significance was determined by one-way ANOVA with Tukey’s multiple-comparisons test. Exact P values are indicated on the graphs. Treatment groups were: (1) YiaT181-Nbs + OmpA-mDR18, (2) YiaT181-Nbs, (3) OmpA-mDR18, (4) commercially available blocking antibodies (αCTLA-4 (clone 9D9) + αPD-L1 (clone 10F.9G2)), (5) OmpA-mDR18 + αCTLA-4 (clone 9D9) + αPD-L1 (clone 10F.9G2), and (6) bacteria-only control.

## Plasmid and primer lists

**Table S1.** Plasmid list.

| Plasmid | Source |
| --- | --- |
| pDS861-RhaRSP-OmpA | This study |
| pDS861-RhaRSP-OmpA-anti-mCTLA-4 | This study |
| pDS861-RhaRSP-OmpA-anti-mPD-L1 | This study |
| pDS861-RhaRSP-OmpA-mDR18 | Yang et al., 2025 |
| pDS861-RhaRSP-YiaT181 | This study |
| pDS861-RhaRSP-YiaT181-anti-mCTLA-4 | This study |
| pDS861-RhaRSP-YiaT181-anti-mPD-L1 | This study |
| pDSG323-Ptet-Neae | Glass et al., 2018 |
| pDSG323-Ptet-Neae-anti-mCTLA-4 | This study |
| pDSG323-Ptet-Neae-anti-mPD-L1 | This study |
| pHEA | Marín et al., 2010 |
| pHEA-anti-mCTLA-4 | This study |
| pHEA-anti-mPD-L1 | This study |
| pAIDA1 | Jarmander et al., 2012 |
| pAIDA1-anti-mCTLA-4 | This study |
| pAIDA1-anti-mPD-L1 | This study |

**Table S2.**
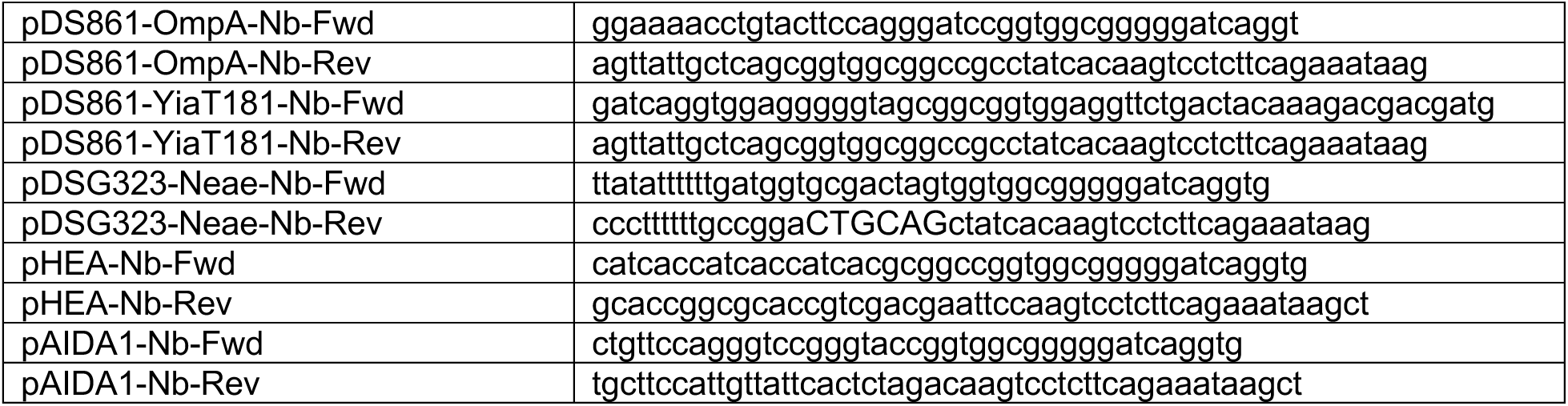
Primers list for cloning.

**Table S3.**
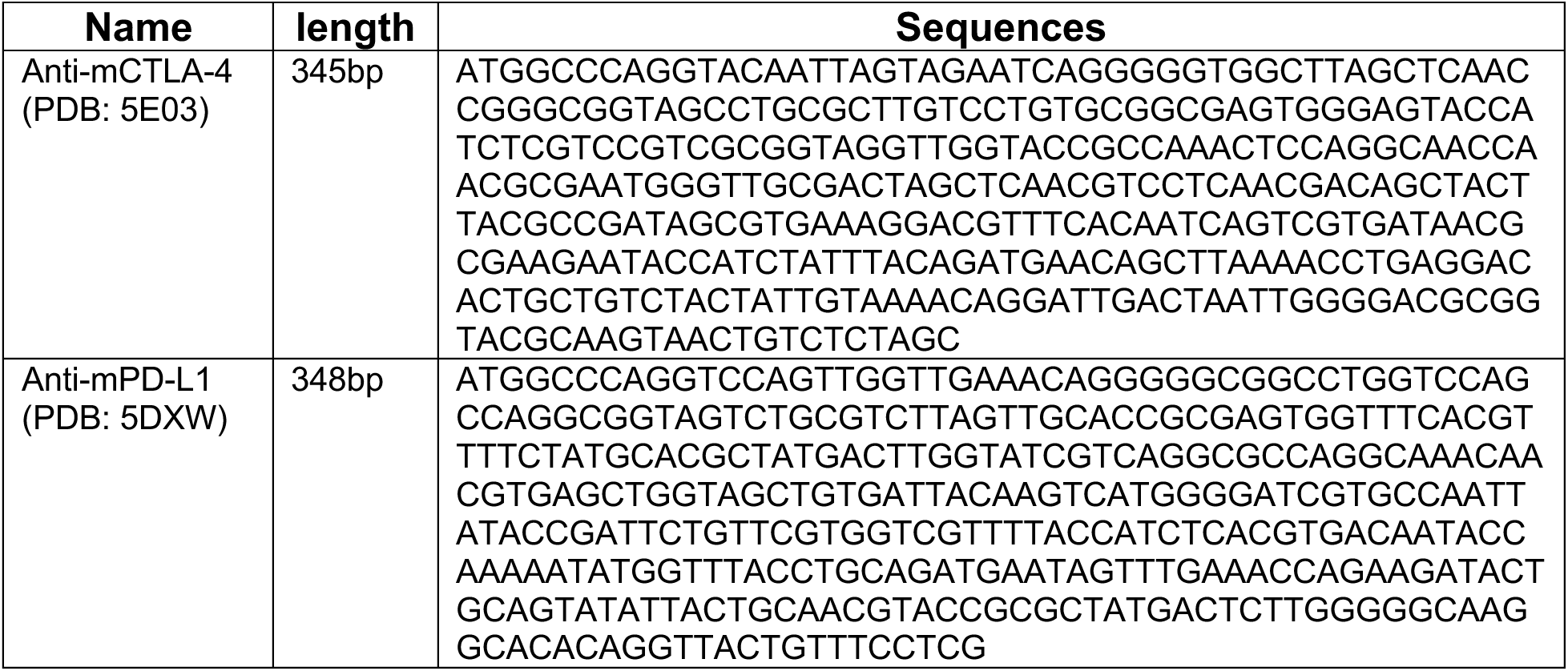
DNA sequence for nanobodies.

**Table S4.** Antibodies used in flow cytometry.

| Target | Dye | Vendor | Catalog | Clone |
| --- | --- | --- | --- | --- |
| Anti-Mouse CD45 | BUV395 | BD Biosciences | 564279 | 30-F11 |
| Anti-Mouse CD8 | BUV496 | BD Biosciences | 569181 | 53-6.7 |
| Anti-Mouse CD4 | BUV615 | BD Biosciences | 613006 | GK1.5 |
| Anti-Mouse PD1 | BUV737 | BD Biosciences | 568362 | J43 |
| Anti-Mouse CD25 | BV421 | BioLegend | 102043 | PC61 |
| Anti-Mouse CD62L | Pacific Blue | BioLegend | 148241 | W18021D |
| Anti-Mouse CD11b | BV480 | BD Biosciences | 566117 | M1/70 |
| Anti-Mouse MHCII | BV510 | BioLegend | 107635 | M5/114.15.2 |
| Anti-Mouse CD86 | BV605 | BioLegend | 105037 | GL-1 |
| Anti-Mouse Ly6C | BV711 | BioLegend | 128037 | HK1.4 |
| Anti-Mouse CD11c | BV785 | BioLegend | 117335 | N418 |
| Anti-Mouse Ly6G | Spark Blue 550 | BioLegend | 127663 | 1A8 |
| Anti-Mouse NK1.1 | PerCP-Cy5.5 | BioLegend | 108728 | PK136 |
| Anti-Mouse Sca1 | RB744 | BD Biosciences | 757749 | E13-161.7 |
| Anti-Mouse TCF1 | PE | Cell Signaling Technology | 14456S | C63D9 |
| Anti-Mouse CD44 | PE-Cy5 | BioLegend | 103009 | 1M7 |
| Anti-Mouse FOXP3 | PE-Cy5.5 | Invitrogen | 15-5773-82 | FJK-16s |
| Anti-Mouse F4/80 | PE-Cy7 | BioLegend | 123113 | BM8 |
| Anti-Mouse CD206 | APC | BioLegend | 141707 | C068C2 |
| Anti-Mouse CD3 | AF700 | BioLegend | 100216 | 17A2 |

